# Early postnatal Flt3^+^ hematopoietic progenitors realize fate-restricted and long-lived output *in vivo*

**DOI:** 10.64898/2026.04.09.716798

**Authors:** Branko Cirovic, Tamar Nizharadze, Nikolaus Dietlein, Carmen Henrich-Kellner, Thomas Höfer, Hans-Reimer Rodewald

## Abstract

Hematopoietic progenitors downstream of hematopoietic stem cells (HSC) are now recognized as the main drivers of day-to-day hematopoiesis. While embryonic and adult HSC fates have been studied in detail, less information exists on stages downstream from HSC, notably in the multipotent progenitor compartment. The early postnatal period represents an important growth phase of the animal and its immune system. Developing immune lineages must be generated in large numbers rapidly, and populate expanding organ niches. To shed light on this critical period, we focused our experiments on early postnatal Flt3^+^ hematopoietic progenitors, and combined genetic single progenitor barcoding using *Polylox* with Flt3-driven, inducible fate mapping. Key immune cell types, including T and B lymphocytes (lymphocytes), innate lymphocytes (ILC) 1-3, NK cells, and granulocytes and monocytes (myeloid) emerged from Flt3^+^ hematopoietic progenitors. Barcode analysis revealed that about 75% of Flt3^+^ hematopoietic progenitors had unipotent fates for lymphocytes, or ILC or myeloid cells, while the remaining fraction showed unprecedented fate combinations for these lineages. Focusing on ILC only, we uncovered clonal fate restriction towards ILC1, or ILC2, or ILC3 in tissues. These data indicate early tissue seeding by progenitors, and further differentiation towards discrete subsets *in situ*. In addition to these fate analyses, induction of fluorescent marker at this intermediate stage of hematopoiesis showed that Flt3^+^ progenitors generated a wave of progeny lasting for over one year. The washout of these cells over time provided kinetic data of cell turnover in major immune cell compartments (in the circulation and in tissues) *in vivo*. In conclusion, we tracked the fate of large numbers (in the order of hundreds) of Flt3^+^ progenitor clones *in situ*. These intermediate progenitors downstream of HSC displayed mostly lineage-restricted fates as well as strong fate complexity, thus serving as a source for early tissue seeding and durable immune lineage.

## Introduction

Hematopoiesis is organized in a hierarchical manner with long-term HSC (LT-HSC) at the top, followed by short-term HSC (ST-HSC), multipotent progenitors (MPP) and downstream lineage-restricted progenitors^1,2^. Our understanding of the function of this system *in vivo* has been advanced in recent years by the development of fate mapping and barcoding experiments^3,4^. These approaches yielded insights into differentiation rates and lineage fates emerging from HSC under native conditions. A key finding from such studies has been a new appreciation of the important roles of progenitors immediately downstream from HSC which appear to have lost self-renewal capacity and initiated differentiation. These heterogeneous progenitors, which can phenotypically be summarized by expression of Flt3^5-7^, contain ST-HSC, MPP, few myeloid progenitors and common lymphoid progenitors (CLP). Originally, largely based on transplant studies, MPP were considered as short-lived transitory stages^6,8^. In contrast, MPP now emerged *in situ* as long-lived progenitors representing the major source of daily hematopoiesis, as well as the key responders under conditions of emergency^9,10^. MPP subsets have been defined phenotypically, and ontogenetically into embryonic and HSC-derived MPP lineages^11,12^. Recent fate mapping experiments revealed that both lymphoid and myeloid lineages derive from MPP *in situ*. However, a comprehensive understanding of the fates across lineages and organs provided by Flt3^+^ progenitors is lacking.

Cell barcoding such as Polylox marks individual cells *in vivo* which can subsequently proliferate and grow into clones^13,14^. Hence, combining the transgenic Flt3-dependent Cre-driver *Flt3creERT2tg*^15^ with Polylox barcoding now enables determination of the native fates, and functions of large numbers of individual Flt3^+^ progenitor clones. Inducible barcoding also offers the possibility to choose the timepoints of interest for label initiation. Most data on hematopoiesis is derived from embryonic (early development) or adult (maintenance) stages. Of note, the early postnatal period represents an important growth phase of the animal in which very large numbers of innate and adaptive immune cells must be generated and must populate growing organs. Additionally, the animal is now directly exposed to environmental stimuli and establishes a gut microbiome^16,17^.

To address the role Flt3^+^ progenitors in this critical postnatal period and subsequently in adult life, we analyzed clonal fates and differentiation dynamics emerging from these progenitors *in situ*. We found that Flt3^+^ progenitors substantially contributed to all major cell lineages including T and B cells, innate lymphocytes (ILC) subsets, natural killer (NK) cells, and myeloid cells (neutrophils and monocytes). Such progeny of Flt3^+^ progenitors was found in all analyzed lymphoid and non-lymphoid organs. Hence, Flt3^+^ progenitors represent a major source to initially build up the immune system. However, the contribution of early postnatal Flt3^+^ progenitors was not only transient but lasted, based on the labeling kinetics in short-lived cells, for at least six months. Barcoding also revealed insights into lineage relationships among progeny from Flt3^+^ progenitors. In contrast to LT-HSC, which yielded predominantly shared barcodes across lineages consistent with multipotency^13^, Flt3^+^ progenitor fates were largely restricted to individual lineages and tissue sites. We also analyzed in detail ILC ontogeny and their lineage relationships, an area that remains poorly understood. Interestingly, most ILC originated from unipotent Flt3^+^ progenitor clones. Tissue analysis combined with *ex vivo* barcoding showed that Flt3^+^ progenitors had seeded tissues by postnatal day six, and generated ILC locally.

In brief, we labelled hundreds of early postnatal Flt3^+^ hematopoietic progenitor cells (HPC) *in situ*, and traced their progeny at clonal resolution. Our results indicate that these progenitors not only build up major compartments in the immune system during the first weeks of life, but also maintain hematopoietic output throughout most of adulthood. At clonal resolution and in contrast to LT-HSC, Flt3^+^ progenitors displayed pronounced fate restriction towards major hematopoietic lineages. Overall, we provide a comprehensive landscape of fates and differentiation rates towards major leukocyte fractions arising immediately downstream of LT-HSC *in vivo*.

## Results

### Barcode initiation in early postnatal Flt3^+^ HPC downstream of HSC

We barcoded Flt3^+^ progenitors using *Polylox*^13^ on day 6 after birth (P6), a time point when organ development is largely completed, yet seeding and filling of peripheral niches by immune cells is an ongoing process in the growing animals (Fig. 1A). Hematopoietic cell differentiation into all lineages transits through a Flt3^+^ cell state^7^. Expression of Flt3 (Fms related receptor tyrosine kinase 3), also known as Flk2 or CD135, tightly correlates with loss of self-renewal of HSC^6,8^ and HSC differentiation^5^. Onset of Flt3 expression occurs in short-term HSC and Flt3 expression marks major progenitor subsets (Fig. S1A-D). We tested whether the *Flt3CreERT2tg*^4,15^ inducible Cre-driver faithfully reproduced the expression pattern observed by flow cytometry. Two days after injection of hydroxy-tamoxifen (OHT) in *Flt3CreERT2tgRosa*^*Polylox*^ mice, that contain both the Flt3-dependent Cre-driver and *Polylox* barcoding cassette, recombined *Polylox* DNA patterns were found only downstream from LT-HSC (lineage^-^Sca1^+^ckit^+^ (LSK) CD150^+^CD48^-^) (Fig. 1B). A prerequisite for fate mapping is induction of label in progenitors but not mature cells. In line with this requirement, we found absence of barcode induction in all tested downstream immune subsets, including ILC subsets, NK cells, T and B cell, and myeloid cells in lung and lymph nodes, blood and T cell progenitors in the thymus (Fig. 1B). When barcodes were analyzed 8 weeks after induction in *Flt3CreERT2tgRosa*^*Polylox*^ mice, recombination was found in all lineage, and large fractions of cells were labelled (Fig. 1C). This fate mapping experiment included immune cells (for cell types see Fig. 1C and Fig. S1E-I), in cervical, peripheral (pooled axillary and inguinal) and mesenteric lymph nodes, lung parenchyma, small intestine lamina propria and intraepithelial lymphocyte fraction, peritoneal lavage, salivary gland, blood, spleen, and thymus. Hence, postnatal day 6 Flt3^+^ progenitors generate within 8 weeks all lineages and cell types analyzed. On these grounds, *Polylox* barcoding in *Flt3CreERT2tg* mice is useful to resolve the underlying fates that generate immune compartments in this critical early phase of life.

**Fig. 1.**
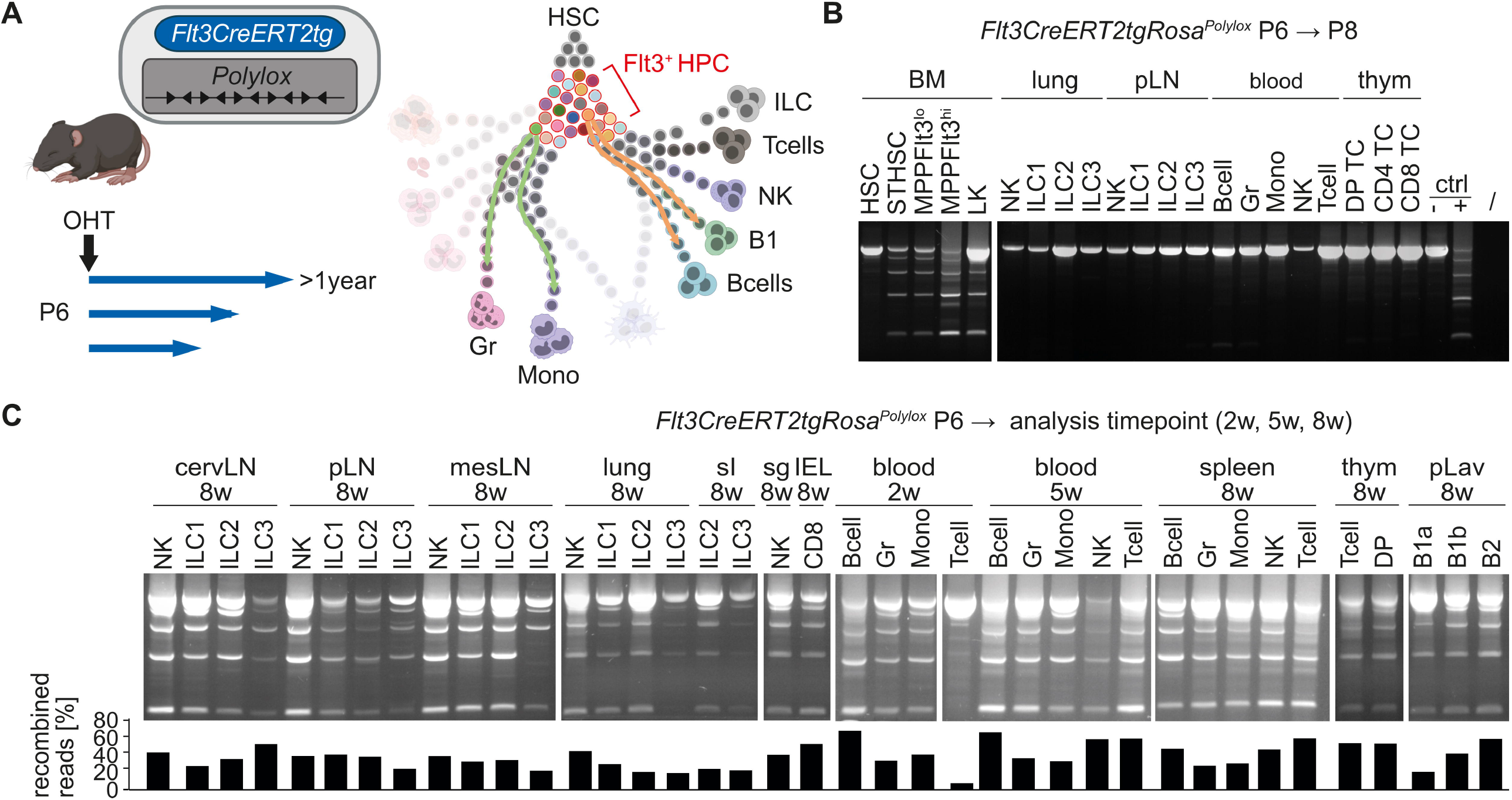
Barcode initiation in early postnatal Flt3^+^ HPC downstream of HSC. (A) Schematic overview of the experimental setup to barcode and trace early postnatal Flt3^+^ HPC. (B) Short OHT-pulse experiment to assess initial barcode placement in *Flt3CreERT2tgRosa*^*Polylox*^ mice. OHT was administered via one single intraperitoneal (i.p.) injection and progenitor cells from bone marrow and immune subsets from the periphery were sorted after two days to assess excision events by DNA amplification of the barcode cassette using flanking primers. BM, bone marrow; Gr, granulocytes, NK, natural killer cells; LK, lineage negative ckit positive cells; DP TC, CD4 CD8 double positive T cells; ctrl, control cell lines either treated with OHT (‘+’, positive control) or left untreated (‘-’, negative control); /, no template control. (C) Top, amplification of the barcode cassette of sorted immune cell subsets from a *Flt3CreERT2tgRosa*^*Polylox*^ mouse injected once with OHT at P6 and analysed two (2w), five (5w) weeks based on blood biopsies and eight weeks (8w) as endpoint after organ harvest. Bottom, histograms indicate the corresponding fraction of barcoding based on the percentage of the reads in non-germline barcode configuration (“recombined reads”). cervLN, cervical lymph nodes; pLN, peripheral lymph nodes; mesLN, mesenteric lymph nodes; sI, small intestine; IEL, intraepithelial lymphocytes (small intestine); thym, thymus; pLav, peritoneal lavage.

### Label propagation dynamics of early postnatal Flt3^+^ HPC

The *PolyloxExpress* locus^18^ expresses tdtomato, and the levels of expression are recombination-dependent. High reporter expression is associated with barcode recombination, while reporter low cells are unrecombined. Hence, in this system, following Cre activity, a fluorescent reporter is permanently expressed which allows quantitative cell tracing by flow cytometry^9^ (Fig. 2A, Fig. S2A). Two days after OHT treatment of *Flt3CreERT2tgRosa*^*PolyloxExpress*^ mice, the fractions of tdtmato^hi^ cells and Flt3^+^ cells (by antibody staining) highly correlated, validating the fluorescent tdtomato marker as live-cell reporter of labeled Flt3^+^ HPC (Fig. 2B). Acute Flt3^+^-dependent Cre activation marked a range of phenotypically defined hematopoietic progenitor subsets downstream of HSC, including early progenitors ST-HSC and MPP, as well as more restricted late progenitors within the Lin^-^Sca1^-^ckit^+^ (LK) fraction, including common lymphoid progenitors (CLP) and to a lesser degree common myeloid progenitors (CMP) and monocyte-granulocyte progenitors (GMP) (Fig. 2B).

**Fig. 2.**
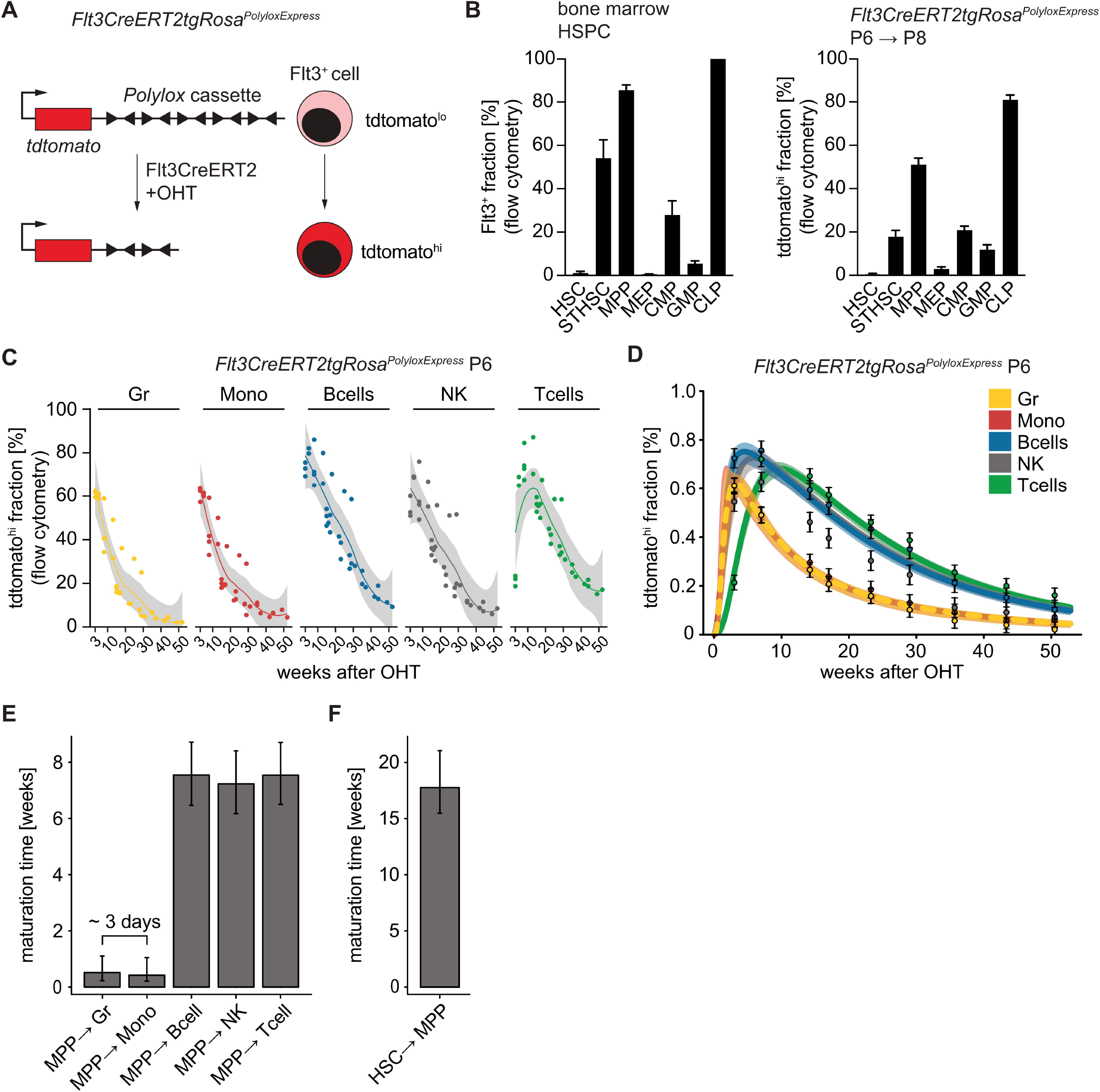
Label propagation dynamics of early postnatal Flt3^+^ HPC. A) Schematic representation of the *PolyloxExpress* barcoding and reporter system. B) Percentage of Flt3-expressing cells across hematopoietic stem and progenitor cell (HSPC) subpopulations in the bone marrow based on flow cytometry (left). Quantification of the tdtomato^hi^ fraction in the corresponding HSPC populations two days after OHT administration in P6 *Flt3CreERT2tgRosa*^*PolyloxExpress*^ mice (right; n=3). C) Kinetic analysis of tdtomato^hi^ cell fractions in peripheral leukocyte subsets in P6-induced *Flt3CreERT2tgRosa*^*PolyloxExpress*^ mice. Serial blood biopsies were taken during overlapping time intervals (3-52 weeks after induction) from six mice in total. Individual data points and smoothing splines (gray) are shown for the indicated chasing duration. D) Model fit of tdtomato^hi^ cell fractions dynamics in peripheral blood for up to 52 weeks post induction in P6-induced *Flt3CreERT2tgRosa*^*PolyloxExpress*^ mice. Data points represent averages of binned tdtomato^hi^ cell fractions. Error bars show standard error of the mean. Curves represent the median fit of the ordinary differential equation compartment model of hematopoiesis fitted to the label dynamics in P6-induced *Flt3CreERT2tgRosa*^*PolyloxExpress*^ mice and previously published data from E9.5-induced *Tie2*^*MCM*^*Rosa*^*YFP*^ mice^21^. Bands around the fitted curves (lighter colors) show 95% confidence intervals. E) Inferred average maturation time of MPPs to granulocytes, monocytes, NK, B and T cells. Maturation time was computed by summing the reciprocals of differentiation rate estimates from MPP to mature leukocytes in the mathematical model in D. Error bars show 95% confidence interval. F) Inferred average maturation time of HSCs to MPP stage. Maturation time was computed by summing the reciprocals of differentiation rate estimates from HSC to MPP in the mathematical model in D. Error bars show 95% confidence interval.

Short term analysis over the first 11 days after OHT induction captured the initial influx of fluorescently labelled cells into leukocyte subsets isolated from peripheral blood (Fig. S2B). Of note, Flt3^+^ HPC rapidly differentiated and progressively contributed to major myeloid and lymphoid lineages, including granulocytes, monocytes, NK cells and B cells, in the periphery. By day 11, all cell types but T cells were found to be labelled in the blood. Owing to their thymic differentiation, labelled T cells appeared in the blood with a delay of 3-5 weeks (Fig. 1C; Fig. 2C). We performed serial bleedings for over one year after induction and quantified the labeling dynamics in distinct cell subsets. The quantitative contribution of Flt3^+^ HPC labeled at P6 to major cell type fractions reached about 60-80% within several weeks after label induction (Fig. 2C). The reporter data is consistent with the quantification of the fraction of recombined reads in *Flt3CreERT2tg*-barcoded mice (Fig.S2C). Hence, at this early postnatal stage, Flt3^+^ HPC are the major source of all analyzed immune cells. The presence of a labelled cell in the periphery can reflect its generation, its life span, or both. For instance, granulocytes are considered to be short-lived (lifespan of several days^19^), and hence their presence in peripheral blood at 30 weeks indicates their recent production from Flt3 fate-mapped progenitors. To gain information on forward differentiation rates from Flt3^+^ HPC, we first analyzed the kinetic data (Fig. 2C), and modeled label propagation from Flt3^+^ progenitors towards mature blood lineages with an ordinary differential equation model based on hematopoietic stem and progenitor cell proliferation and differentiation (Fig. 2D; see mathematical modeling of label dynamics in mature blood populations in Methods). The model captures the generation of the tdtomato^hi^ cell fraction in granulocytes, monocytes, B cells, T cells, and to a lesser extent, NK cells. Differentiation rates from MPP towards myeloid and lymphoid lineages were previously estimated based on label propagation emerging further upstream from Tie2^+^ HCS^20,21^. Measurements of output rates are expected to be more precise when products are closer the source, hence, our Flt3-driven fate mapping now offered an opportunity to scrutinize this key differentiation step from MPP to myeloid and lymphoid lineages kinetically. In line with our previous Tie2 fate mapping results, the step from MPP to lymphoid lineages, now resolved into B, T and NK cells, took on average in the order of 7 weeks (Fig. 2E). In contrast, the generation of both granulocytes and monocytes was very rapid (in the order of three days) (Fig. 2E). From the previous Tie2 fate-mapping we inferred that the transition of MPP to CMP was faster than 2.6 days, without being able to resolve differentiation steps downstream of CMP^21^. Hence the labeling closer to the source indeed gave a more accurate estimate of myeloid differentiation time that, however, remained consistent with Tie2 fate mapping. In summary, consistent differentiation rates from MPP to myeloid and lymphoid lineages were obtained comparing Flt3- or Tie2-driven fate mapping.

Given that label was induced in *Flt3CreERT2tg* mice at an intermediate stage of the hematopoietic tree (Fig. 1A, Fig. 1B), the decreasing tdtomato^hi^ leukocyte fractions with time (Fig. 2D) were due to ‘washout’ from differentiating unlabeled upstream compartments (HSC and ST-HSC)^22,23^. The long timescale of the replacement by unlabeled cells was determined by the slow maturation of HSC into MPP, taking 18 weeks on average (Fig. 2F). Comparison of label propagation rates obtained previously^21^ with the inferred rates from this study using Flt3 fate mapping, found that differentiation rates from HSC to ST-HSC and to MPP were congruent (Fig. S2D). Label retention over time (2 days to 16 weeks) in each stem and progenitor compartment again visualized the rare input from HSC into ST-HSC (label was retained over this period), and the higher replacement of, for example, MPP and CLP from upstream compartments (Fig. S2E). We extended this kinetic analysis beyond the 16 weeks by barcode analysis. While at 22 weeks post label induction a fraction of ST-HSC and progenitors for all lineages still carried barcodes (i.e. remained labelled) (Fig. S2F, bottom), by 53 weeks, the abundance of barcodes was strongly reduced (Fig. S2G, bottom). Based on reporter analysis at this time point, percentages of tdtomato^hi^ leukocytes ranged from 2-17% in the blood (Fig. 2C). In summary, these data show that Flt3^+^ HPC at P6 contribute long-term to the generation of diverse immune cells for at least one year, which is consistent with earlier measurements showing longevity of MPP^24-26^. Remarkably, differentiation rates from HSC to downstream ST-HSC and onwards to progenitors were similar regardless of label induction in HSC via *Tie2-* or at intermediate stages via *Flt3*-driven fate mapping.

### Clonal Flt3^+^ HPC fate combinations towards major leukocyte lineages

To measure fates or fate combinations emerging from early postnatal Flt3^+^ HPCs, we induced barcodes in *Flt3CreERT2tgRosa*^*Polylox*^ mice at P6, and analyzed barcodes after 22 weeks in immune subsets in a wide range of lymphoid and non-lymphoid tissues and blood (Fig. 1A, right; Fig. S2F). Across all cell types and lineages, we detected 1652 distinct barcodes. A subset of barcodes will be generated with high probability and hence in more than one progenitor cell; to eliminate these redundant barcodes (which might be mistaken as fate sharing), we filtered for barcodes with low probability of generation (*P*_gen_ < 10^−4^), resulting in 976 barcodes for further analysis (Fig. 3A)^14^. Indeed, barcodes with *P*_gen_ < 10^−4^ are in the frequency plateau characteristic of clonal barcodes^49^ (Fig. S3A), show stable proportions of multilineage and unilineage fates (Fig. S3B), while retaining about two thirds of all detected barcodes (Fig. S3C). Collectively, we generated and analysed a large repertoire in the range of ∼1000 distinct, clonal barcodes from an induced *Flt3CreERT2tgRosa*^*Polylox*^ mouse (Fig. 3A). For comparison, we induced barcodes in *Tie2*^*MCM*^*Rosa*^*Polylox*^ mice^13,21^ on embryonic (E) day 9.5, and analyzed the HSC fates at adult age (Fig. 3B). In both cases, *Flt3CreERT2tgRosa*^*Polylox*^ and *Tie2*^*MCM*^*Rosa*^*Polylox*^ mice, we comprehensively sampled progenitors in bone marrow and thymus, and in the periphery ILC subsets 1-3, NK cells, T cells, and B1 and B2 cells, granulocytes and monocytes in, where appropriate, cervical, peripheral (pooled axilliary and inguinal) and mesenteric lymph nodes, lung parenchyma, small intestine lamina propria and intraepithelial lymphocyte fraction, peritoneal lavage, salivary gland, blood, spleen, and thymus (Fig. 3A, B).

**Fig. 3.**
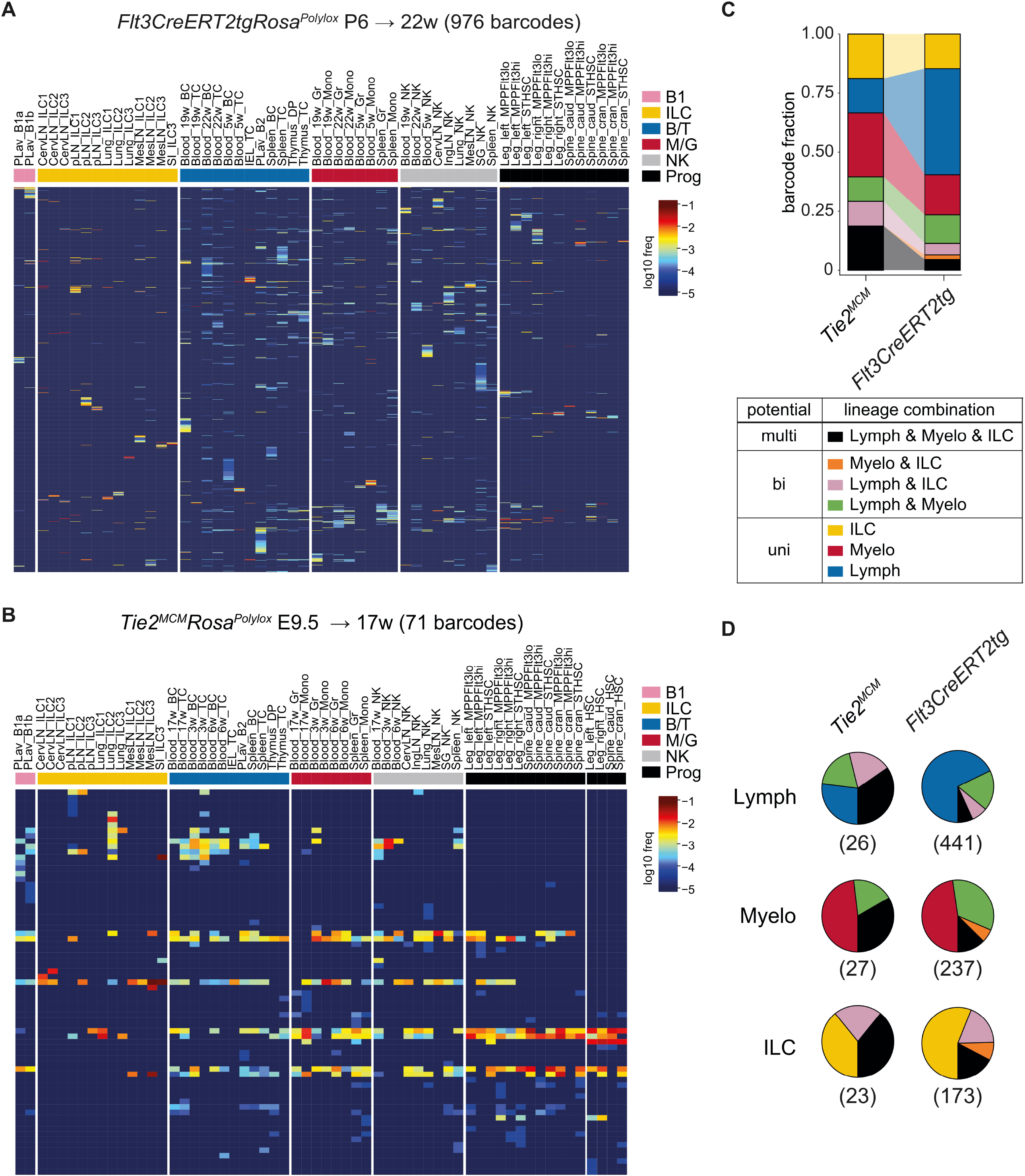
Clonal Flt3^+^ HPC fate combination towards major leukocyte lineages. A) Barcode landscape derived from a *Flt3CreERT2tgRosa*^*Polylox*^ animal induced at P6 and analysed 22 weeks later (final analysis timepoint). Heatmap shows the frequency of 976 barcodes (in rows, filtered for *P*_gen_ < 10E-4) across cell subsets and organs (in columns). Cell populations are organised in color-coded groups. B) Corresponding result from a *Tie2*^*MCM*^*Rosa*^*Polylox*^ animal induced at E9.5 and analysed 17 weeks later (final analysis timepoint). C) Comparison of barcode sharing across cell lineages for barcoded P6 Flt3^+^ HPC (22 weeks after induction, (as in A) and E9.5 barcoded Tie2^+^ HSC (17 weeks after induction, as in B). Proportions of barcodes found exclusively in one leukocyte class or shared between classes are respectively color-coded (see legend) and categorized as ‘uni-’, ‘bi-’ or ‘multi-’potent. D) Pie charts indicating lineage-centric views focusing on clonal barcodes pre-filtered according to the leukocyte classes “Lymph” (B and T cells), “Myelo” (monocytes and granulocytes) and “ILC” (helper ILC1-3). Number of included barcodes are shown in brackets. Color annotation as in C.

Visual inspection of the heatmaps showing the resulting barcode patterns suggested mostly non-shared barcodes in *Flt3CreERT2tgRosa*^*Polylox*^ mice (Fig. 3A) as opposed to mostly shared barcodes in *Tie2*^*MCM*^*Rosa*^*Polylox*^ mice (Fig. 3B) (fate combinations summarized in Fig. 3C). To test these data versus a null model, in which only single fates occur, we quantified fates emerging after *Flt3CreERT2tg*- or *Tie2*^*MCM*^- driven barcoding (Fig. S3D). Indeed, greater barcode sharing was seen in *Tie2*^*MCM*^*Rosa*^*Polylox*^ compared to *Flt3CreERT2tgRosa*^*Polylox*^ mice, and, moreover, a mathematical model of barcode generation in HSC or MPP and subsequent propagation yielded the same result (Fig. S3E). In simple terms, these data would be consistent with the general idea of progressive lineage commitment with hematopoietic differentiation.

We classified barcodes across organs and cell types for their presence or absence in “Lymph”, consisting of conventional B2 and T cells, in “Myelo”, consisting of granulocytes and monocytes, or in “ILC”, consisting of a combination of the helper ILC subsets ILC1-3. This analysis disregards further separation within each group (for example B versus T cells). Barcodes detected in one of the three leukocyte classes are termed “unipotent”, in two “bipotent”, and in all three classes “multipotent” (Fig. 3C). The overall fates according to this classification were enriched for multilineage in *Tie2*^*MCM*^*Rosa*^*Polylox*^ mice, and for unipotent in *Flt3CreERT2tgRosa*^*Polylox*^ mice (Fig. 3C). These analyses reflect the fates emerging from a given stem or progenitor cell (‘onward looking’). To address fates from the view of a target compartment, we introduced ‘lineage centric views’, by asking which fates contributed to what extent to the formation of a mature compartment. Lymph-, Myelo-, or ILC-centric views were compared for *Flt3CreERT2tgRosa*^*Polylox*^ and *Tie2*^*MCM*^*Rosa*^*Polylox*^ mice (Fig. 3D). For instance, analysis of *Flt3CreERT2tgRosa*^*Polylox*^ mice showed that the Lymph compartment is filled to about 66% from Lymph unipotent fates (blue area), with smaller contributions from bipotent Lymph-Myelo-, or Lymph-ILC-, and from multi-potent fates. Similarly, in the order of 50% unipotent fates generate the Myelo (red) and ILC (yellow) compartments (Fig. 3D). Hence, mature immune cell compartments are generated from progenitors exhibiting different fates, with major contributions from lineage-restricted progenitors.

While these analyses were based on barcode abundance, we next also considered progenitor clone sizes. Barcode read counts can serve as a proxy for clone sizes^24^. Interestingly, for both *Flt3CreERT2tgRosa*^*Polylox*^ and *Tie2*^*MCM*^*Rosa*^*Polylox*^ data, when considering progenitor clone sizes, the contributions to all compartments of multi-potent fate clones were larger (Fig. S3F) compared to the barcode contributions based on barcode presence alone (Fig. 3D). This was in particular pronounced in the *Tie2*^*MCM*^*Rosa*^*Polylox*^ data (Fig. 3D). These findings indicate that larger clones tend to possess more multilineage fates, which, in turn, suggests that stronger clonal expansion is not associated with fate loss. Finally, the observed fate combinations were similar at all analysis timepoints, ranging from two months until over one year after barcode induction (Fig. S3G).

### Subset- and tissue-resolved fate specification of early postnatally-labeled Flt3^+^ HPC

The rich data sets from *Flt3CreERT2tg* barcoded mice, offers a deeper view into the developmental relationship of lymphocyte subsets arising from Flt3^+^ progenitors. Flt3^+^ MPP and CLP have both been invoked as key stages in lymphocyte development, however, the fates of individual Flt3^+^ progenitors remain unknown. In particular, the origins and relatedness of ILC to other lymphocytes has been subject to debate^27-30^. We first compared barcodes that are present in innate-like B1a/b cells (“B1”), helper ILC1-3 (“ILC”), conventional T cells (“T”) (Fig. 4A) or conventional B cells (“B”) (Fig. S4A). Applying this filtering and comparison approach, we observed a majority of detected barcodes induced in single Flt3^+^ progenitor cells exclusively in one individual lymphocyte subgroup and only marginally shared among each other (Fig. 4A, left top). By contrast, a considerable fraction of barcodes was shared among the corresponding subgroups in *Tie2*^*MCM*^-induced mice (Fig. 4A, left bottom). Further resolution of helper ILC subsets into ILC1-3 similarly revealed minor overlap between ILC subsets in *Flt3CreERT2tg* mice, suggesting that most Flt3^+^ progenitors had predetermined ILC subset fates (Fig. 4A, middle).

**Fig. 4.**
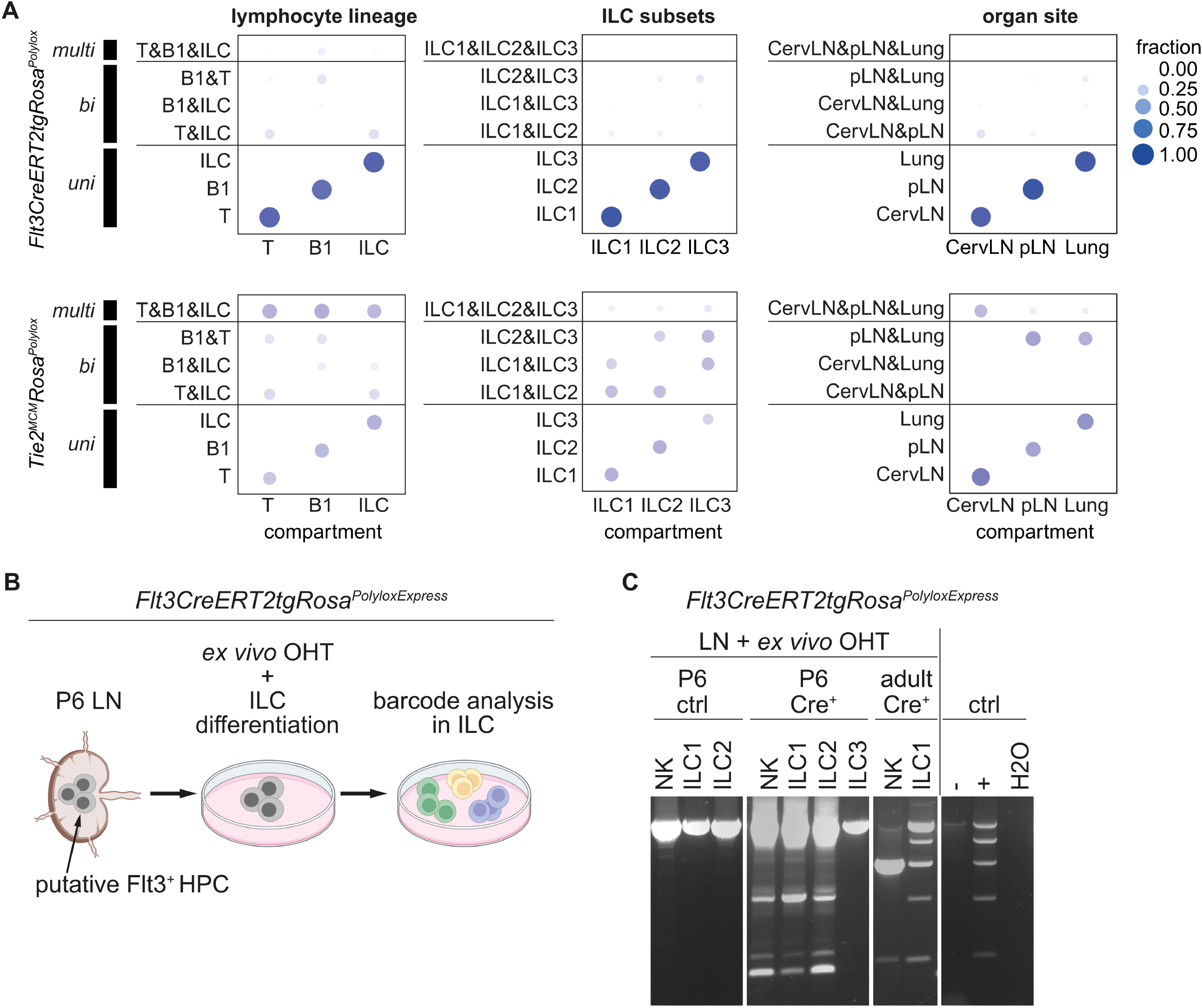
Subset- and tissue-resolved fate specification of P6-labelled Flt3^+^ HPC clones. A) Subtype- or organ-centric barcode sharing based on *Flt3CreERT2tgRosa*^*Polylox*^ (top sections) or *Tie2*^*MCM*^*Rosa*^*Polylox*^ OHT-induced animals (bottom section) that are presented in Figure 3. Individual columns of the bubble plots represent barcodes filtered according to the respective indicated compartments of interest. Bubble size and shading in rows indicate the respective barcode fractions contributing to individual fate subsets (“uni”), combination of two subets (“bi”) or all three subsets (“multi”). Comparisons are summarised for lymphocyte lineages (left), ILC subsets (middle) and organ sites (right). B) Experimental approach for *ex vivo* barcode induction in Flt3^+^ HPCs derived from LN suspensions and subsequent *in vitro* ILC differentiation and analysis. C) Barcode amplification from *in vitro* differentiated and purified ILC subtypes (*Flt3CreERT2tgRosa*^*PolyloxExpress*^ mouse). Lymph nodes from a Cre negative animal (P6 ctrl) served as control. Lymph nodes were isolated either from P6 animals or adult (>8 weeks) mice. Ctrl, control cell lines either treated with OHT (‘+’, positive control) or left untreated (‘-’, negative control); H2O, no template control.

Given the tissue residency of ILC^31^, and evidence for the presence of ILC progenitors in tissues^31-33^, we next focused on ILC subsets in tissues (lung, cervical lymph node and pool of inguinal and axillary lymph nodes (pLN)). Strikingly, we found little evidence for common Flt3^+^ progenitors generating ILCs in the sampled tissues (Fig. 4A, right). Progenitor differentiation within tissues may underlie these findings. However, a confounding effect might be that technical undersampling could make barcodes arising from a common progenitor appear tissue-restricted. To test for this, we simulated *in silico* that all barcodes arose from common progenitors for ILC1, ILC2 and ILC3 (Fig. S4B, middle panel), and applied the effect of experimental sampling (Fig. S4B, right panel). We conclude from this simulation that subsampling would preserve barcode origins from common precursors. Indeed, sampling as little as 10% of ILCs would still reveal common progenitors by barcode sharing (Fig. S4C). Hence, the distinct barcodes found in different locations are not due to undersampling. Taken together, the data suggest that the ILC fates of Flt3^+^ progenitor clones are already separated at day six after birth and thus distinct barcodes are inherited by the ILC subpopulations, and ILC in tissues.

### Flt3^+^ HPC in postnatal lymph nodes generate ILC locally

Next, we asked whether labelled ILC progeny originated from central Flt3^+^ progenitors (bone marrow) or from peripheral tissue-resident Flt3^+^ progenitors. We devised an experiment to identify Flt3^+^ progenitors in lymph nodes of P6-old *Flt3CreERT2tgRosa*^*PolyloxExpress*^ animals (Fig. 4B). In this setup, short *ex vivo* OHT treatment of the lymph node cell suspension targets putative Flt3^+^ progenitors *in situ*, and excludes central labelling or direct labeling of mature (Flt3^-^) ILCs (Fig. 1B). After three weeks of culture supporting ILC differentiation^30^, cells belonging to all ILC subsets as well as NK cells were detected by flow cytometry sorting into ILC1-3, and following barcode analysis of the subsets (Fig. 4C). Importantly, all but the ILC3 subset contained recombined barcodes, demonstrating Flt3^+^ progenitors in P6 lymph nodes for ILC1 and ILC2. The potential for generation of ILC1 and NK cells was retained in adult lymph nodes. Here, ILC2 and ILC3, which were present in the lymph nodes *ex vivo* (Fig. 1B, C), were undetectable post culture. In summary, early in life, Flt3^+^ progenitors are found in lymph node and possibly other peripheral organs where they directly give rise to specific immune subsets *in situ*. The relative contributions of central versus peripheral progenitors for the full generation of ILC compartments *in vivo* remains to be determined.

## Discussion

Direct fate comparison between HSC and Flt3^+^ progenitors revealed marked differences. As expected, HSC had abundant multilineage fates. Based on barcode reads, which is proxy for progeny clone size, at least 75% of HSC clones generated lymphocytes, myeloid cells and, notably, also ILC (Fig. S3F). Using the Flt3-driver for barcoding, we initially labelled a mix of ST-HSC, MPP, myeloid progenitors and CLP in the proportions shown in Fig. 2B. While it is difficult to directly assign fates to each of these phenotypically defined populations, we note that overall, only in the order of 25% Flt3^+^ progenitors had multilineage fates for lymphocytes, myeloid cells and ILC (Fig. 3C). This indicates that the onset of major fate restrictions coincides with expression of Flt3. These fate restrictions ranged from single lineage fates (lymphocytes only, myeloid cells only, or ILC only) to combined fates for two but not all lineages (e.g. lymphocytes and ILC but no myeloid cells). Regarding ILC, most ILC were generated from ILC-restricted fates, followed by shared ILC-lymphocyte fates and, more rarely, from shared ILC-myeloid fates. This fate analysis appears incompatible with a simply annotation of ILC as ‘lymphoid’, at least based on ontogeny. Zooming in on lineage relationships between lymphocytes (T, B1, B2) and ILC, demonstrated mostly uni-lineage fates for each on these lineages (Fig. 4A, Fig. S4A). Interestingly, also the ILC1-3 subsets showed little fate overlap with each other (Fig. 4A), and ILCs residing in different organs were unrelated with each other based on fates. The latter findings suggest early seeding of tissues with Flt3^+^ progenitors which were then barcoded on postnatal day 6 *in situ*. This model is strongly supported by the *ex vivo* labeling of Flt3^+^ lymph node cells and their subsequent *in vitro* differentiation into ILC.

While progenitor differentiation has generally been considered to take place centrally in the bone marrow, our data, together with previous findings, including development of ILC1 in the liver ^33^ and ILC2 in the lung ^32^, indicates that tissue-resident cells can be generated from local progenitors. Regarding ILC development within hematopoiesis, models suggest that ILCs arise from CLP via several intermediate stages (common ILC progenitor (CILCP), common helper-ILC progenitor (CHILP) and distinct ILC precursors (ILCp))^34^. Interestingly, entry into this differentiation pathway towards ILC is associated with downregulation and loss of Flt3 expression (high in CLP, CHILP and lost in ILCp)^35,36^. Moreover, the terms CILCP and CHILP implied a common progenitor for ILC1-3 subtypes, i.e. a single progenitor cell giving rise to multiple ILC lineages. However, the fact that our Flt3-driven barcoding resulted in unique barcodes in either ILC1, ILC2 or ILC3, and no shared barcodes between all these subtypes (Fig. 4A, middle panel) argues against common ILC progenitors in the Flt3-positive compartment and further below. Furthermore, commitment towards ILC1, ILC2 or ILC3 must occur while progenitors are still expressing Flt3. In brief, our data appear incompatible with known models of ILC development.

Despite the observed fate restrictions, Flt3^+^ progenitors were a major source of all analyzed hematopoietic lineages. In fact, comparison of initial labeling frequencies (Fig. 2B, right panel) to peak output (Fig. 2C) indicated roughly a direct proportionality, suggesting that essentially all hematopoietic cells passed through a Flt3^+^ progenitor stage^7^. The presence of labeled granulocytes and thymus progenitors (both short-lived cells) for up to one year after labeling confirms earlier reports of long-lasting hematopoiesis emerging from early postnatal MPP^12,24,26,37^. After the maximum of progeny labeling, frequencies declined in all lineages which is the result of the slow replacement of labeled by unlabeled progenitors (Fig. 2C, 2D). The dynamics of hematopoietic differentiation has been measured previously based on HSC output and label propagation along progenitor stages^21^. This quantitative framework can now be probed based on the washout kinetics of granulocytes, monocytes, B cells, NK cells and T cells following labeling of Flt3^+^ progenitors (Fig. 2C). Interestingly, kinetic modeling showed very rapid differentiation from MPP to granulocytes and monocytes (in the order of three days). In contrast, it took in the order of seven weeks to B cells, NK cells or T cells from MPP, providing independent evidence for previously estimated rates at the entry into myeloid or lymphoid differentiation^21^. Moreover, the kinetics of label decay also provided information on differentiation kinetics upstream of Flt3^+^ progenitor cells. This analysis indicated that it took in the order of 17 weeks to generate MPP from HSC, which is in line with earlier output measurements^21^. Of note, the new HSC output rate has now been obtained independently from a chosen HSC Cre-driver.

In summary, we deconvoluted the Flt3^+^ progenitor compartment into hundreds of individual progenitors and examined their clonal fates under physiological conditions *in vivo*. We found complex clonally restricted fates in this compartment downstream of HSC. The Flt3^+^ progenitor compartment is a major and durable contributor to steady state hematopoiesis. Unexpectedly, the Flt3^+^ progenitor compartment harbors almost exclusively uni-ILC-lineage fates rather than common ILC fates. These experiments indicate commitment for the ILC subtypes prior to the previously described ILC progenitors.

## Methods

### Mice

The barcoding mice *B6-Gt(ROSA)26Sor*^*tm1(Polylox)Hrr*^ (termed *Polylox*) and *B6-Gt(ROSA)26Sor*^*tm1(PolyloxExpress)Hrr*^ (termed *PolyloxExpress*) were generated as described previously^13,18^. Male or female mice derived from crossings of heterozygous *Tek*^*tm1*.*1(icre/Esr1*)Hrr*^ (termed *Tie2*^*MCM*^)^21^ or *Tg(Flt3-cre/ERT2)#Ccb* (termed *Flt3CreERT2tg*)^15^ with barcoding mice were used as experimental animals as indicated. Correct genotypes were confirmed by standard PCR protocols. Mice were kept in individually-ventilated cages under standardized and specific pathogen-free conditions in the animal facility at the DKFZ. All animal experiments were performed in accordance with institutional and governmental regulations, and were approved by the Regierungspraesidium (Karlsruhe, Germany).

### Induction of barcode recombination

Recombination of the barcode cassette *in vivo* was induced by injection with (Z)-4-Hydroxytamoxifen (OHT, Sigma). OHT was first dissolved in pure ethanol at 100mg/ml, heated (55 °C) and vortexed before further dilution (10mg/ml) in pre-warmed peanut oil (Sigma). Aliquots were stored at -20 °C and pre-warmed to body temperature before injection. OHT was used at a concentration of 45 µg per g body weight. P6 old mice were injected intra-peritoneally (i.p.). For induction *in utero*, OHT was supplemented by 22.5 µg per g body weight progesterone (Sigma) and administered by oral gavage. Appearance of vaginal plugs was defined as embryonic day 0.5 (E0.5). Embryonically induced litters were delivered by cesarean section and raised by foster animals to reduce the chance of birth complications.

### Organ preparation and cell purification

For serial blood collection, the submandibular vain was punctured using a lancet. For endpoint analysis mice were euthanized and blood and peritoneal lavage was collected using FACS buffer (5 % FBS, 50 mM EDTA, PBS) before transcardial perfusion was performed with 20 ml cold PBS. Lung, spleen, small intestine, salivary gland, thymus, axillary and inguinal lymph nodes (pooled as peripheral lymph node sample (pLN)), cervical lymph nodes and mesenteric lymph nodes were isolated and stored in ice-cold FACS buffer until further processing. To purify the lamina propria leukocyte fraction from small intestine, fecal material and Peyer’s patches were first removed and tissue cut into smaller pieces. After incubation in 2 % FBS, 5 mM ETDA, 1 mM DTT (in HBSS) for 15 min at 37°C, 200 rpm shaking, the reaction was stopped with 10% FBS, 20 mM HEPES RMPI1640 medium. The material was applied on a 100 µm strained and the flowthrough was collected as intraepithelial leukocyte (IEL) fraction. Organ pieces were further digested in media (2 % FBS, 20 mM HEPES in RPMI1640) supplemented with 50 µg/ml DNaseI (Sigma #DN25) and 200 µg/ml collagenase IV (Sigma, #C5138) for 30 min at 37 °C, 200 rpm. Minced salivary gland and lung were digested accordingly. Digestion was stopped with 10 % FBS, 2 mM EDTA, 20 mM HEPES RMPI1640 medium. Cell suspensions were further purified by filtering and washing through 100 µm and subsequently 40 µm strainers. Spleen, thymus and lymph nodes were minced and directly filtered through a 40 µm strainer. Red blood cells were removed from spleen, lung and blood cell suspensions by incubation with RBC lysis buffer (BioLegend) for 3 min at room temperature. Lymphocyte fractions in lung, IEL, salivary gland and small intestine samples were further enriched by addition of a Percoll gradient centrifugation step (67 % and 44 % double Percoll gradient, centrifugation at 1600 g for 15 min without brake at 4 °C). Hematopoietic progenitors were isolated from the bone marrow by crushing femur and tibia. For more detailed analysis progenitors were processed separately from femur and tibia (pooled and termed ‘hind leg’), humerus (termed ‘front leg’) and vertebral column. The vertebral column was again divided into a cranial part (cervical and thoracic section) and caudal part (caudal and lumbar section). The hematopoietic progenitor cell fraction was further enriched by magnetic bead-based depletion of cells bound by biotinylated antibodies (against CD3e, CD4, CD8a, CD11b, CD19, Gr-1, TER119) following the manufacturer’s guidelines (Miltenyi Biotec).

### Flow Cytometry

Cell suspensions for flow cytometric analysis and sorting were further processed in FACS buffer (5 % FBS, 50 mM EDTA, PBS). To reduce unspecific binding of fluorescently labeled antibodies, cell suspensions were incubated in 1:25 unconjugated mouse IgG (ChromPure, Jackson ImmunoResearch) for 15 min on ice. Cell staining with cocktails of fluorophore-conjugated antibodies (supplemental information) were performed in FACS buffer for 1h on ice in the dark. Combinations of antibodies were used to define specific cell populations as described in the supplementary figure (Fig. S1). After incubation, cells were washed, resuspended in SytoxBlue (Invitrogen, 1:10,000 in FACS buffer) for live/dead discrimination and filtered (40 µm strainer) directly before analysis and sorting. Samples were acquired using BD FACS ARIAIII (BD Biosciences) and the FACS Diva software (BD Biosciences, USA). Data was further analyzed using FlowJo (BD Biosciences, USA). For cell sorting, selected cell populations were collected in filtered 10 % FBS, PBS. Correct sorting conditions were calibrated using Accudrop beads (BD Biosciences, USA) and verified by re-analysis of a fraction of sorted cell-populations.

### ILC differentiation in vitro

5.0E+4 OP9-DL4 feeder cells were seeded the day before induction of ILC differentiation in complete IMDM (cIMDM) medium (IMDM, 10 % FBS, 1 % penicillin/streptomycin, 1 % sodium pyruvate) in a 24-well plate format and incubated over night. Feeder cells were then irradiated (30 Gy; Cesium 137 GammaCell40 Irradiator (Best Theratronics)). Adult or P6 old *Flt3CreERT2tgRosa*^*PolyloxExpress*^ animals were sacrificed under sterile conditions to isolate axillary, subiliac and mesenteric lymph nodes (pooled per animal (pLN)). Bone marrow (BM) was prepared by crushing femur and tibia. Cell suspensions were prepared by gentle filtering through a 40 µm cell strainer and washed in cIMDM medium. 1.0E+5 pLN or BM cells were added to the feeder cells in supplemented cIMDM (cIMDM with 20 ng/ml IL-7 (Miltenyi Biotec), 20 ng/ml SCF (Miltenyi Biotec) and 50 µM betamercaptoethanol). Barcode recombination was initiated *ex vivo* by addition of OHT (100 nM). After 24h medium was carefully replaced with fresh supplemented cIMDM without OH-TAM. Half of the medium was replenished every 2-3 days. Cells were harvested after 3 weeks by trypsinisation and further processed for flow cytometric analysis as described previously.

### Barcode amplification and long-read sequencing

Barcode detection by long-read sequencing (Pacific Biosciences) was essentially done as described previously in detail^14^. Briefly, enriched cell fractions were pelleted at 500 g for 5 min at 4 °C and resuspended in 25 µl cell lysis buffer (1x PCR Buffer 1 from Expand Long Template PCR System (Roche), 0.5 mg/ml Proteinase K (ThermoFisher)). After lysis for 60 min at 56 °C, Proteinase K was inactivated at 95 °C for 10 min. Cell lysates were stored at -20 °C until further processing. Samples were subjected to polymerase chain reaction in a total volume of 50 µl 1x PCR Buffer 1 and Polymerase from Expand Long Template PCR System (Roche) and dNTPs mix (250 nM each; ThermoFisher) to amplify the *Polylox* barcode cassette. Primer pairs #2426 (5’-CGACGACACTGCCAAAGATTTC-3’) and #2427 (5’- CATACCTTAGAGAAAGCCTGTCGAG-3’) (200 nM final concentration for each) were added based on sequences complementary to flanking regions of the *Polylox* cassette. The primer pair sequences were designed to extend in 5’ direction by unique 8-mer barcode handles and universal 5-mer pad sequences (GGTAG) to allow unique combinatorial use for subsequent pooling of PCR amplicons and multiplexed sequencing library preparations (modified #2426 and #2427 resulting in FW_2426_A-X and RV_2427_1-24, respectively; supplemental information). PCR cycling conditions were set as following: 5 min at 95 °C; 35 cycles of 30 s at 95 °C, 30 s at 54 C, 5 min at 72 °C) and final extension step for 10 min at 72 °C. Barcoding efficiency was verified by standard agarose gel electrophoresis. Multiplexed SMRTbell libraries starting from 150 ng DNA input were prepared following the ‘Preparing SMRTbellTM Libraries using PacBio Barcoded Universal Primers for Multiplex SMRT Sequencing’ procedure guidelines (Pacific Biosciences) and using reagents from ‘SMRTbell Express Template Prep kit 2.0’ and Sequel sequencing kit 3.0 (Pacific Biosciences). Libraries were sequenced on SMRT Cell 1M using the SMRT Link interface and Sequel System (Pacific Biosciences).

### Pre-processing and downstream analysis of barcode sequencing reads

The used computation pipeline for barcode retrieval and evaluation from raw long-read sequencing data was described previously^14^. Pre-processing of raw sequencing data was performed within the SMRT Link software suite (Pacific Biosciences). Circular Consensus Sequencing (CCS) reads were called using the following parameters: Minimum number of passes=1; minimum predicted accuracy=0.9; minimum CCS read length=200; maximum CCS read length=2400. Demultiplexing of samples was done with the lima package (Pacific Biosciences) and matching of both forward and reverse barcoded primers (8-mer + #2,426 or #2,427) was used to robustly call sample identities. Barcode calling was performed following the RPBPBR pipeline (https://github.com/hoefer-lab/RPBPBR.git) with default settings that includes 5’/3’-end trimming, alignment of barcode segments and subsequent barcode assembly. Probabilities of barcode generation (P_gen_) were calculated as previously described (https://github.com/hoefer-lab/polylox.git) and a threshold was set to 1.0E-4 to remove uninformative barcodes for further analysis. Data visualization of barcode frequencies was done within the RStudio environment using standard software packages (gglot2, pheatmap, complexheatmap).

### Mathematical modeling of label dynamics in mature blood populations

The dynamics of tdtomato^hi^ cell fraction in blood granulocytes, monocytes, NK cells, B cells and T cells were modeled with an ordinary differential equation model. The model contained following stem and progenitor populations: HSC, ST-HSC, MPP, CMP, CLP, GMP, as well as mature populations granulocytes (Gr), monocytes (Mono), B cells (B), NK cells (NK), T cells (T) and their respective maturation stages. Model equations are as follows:

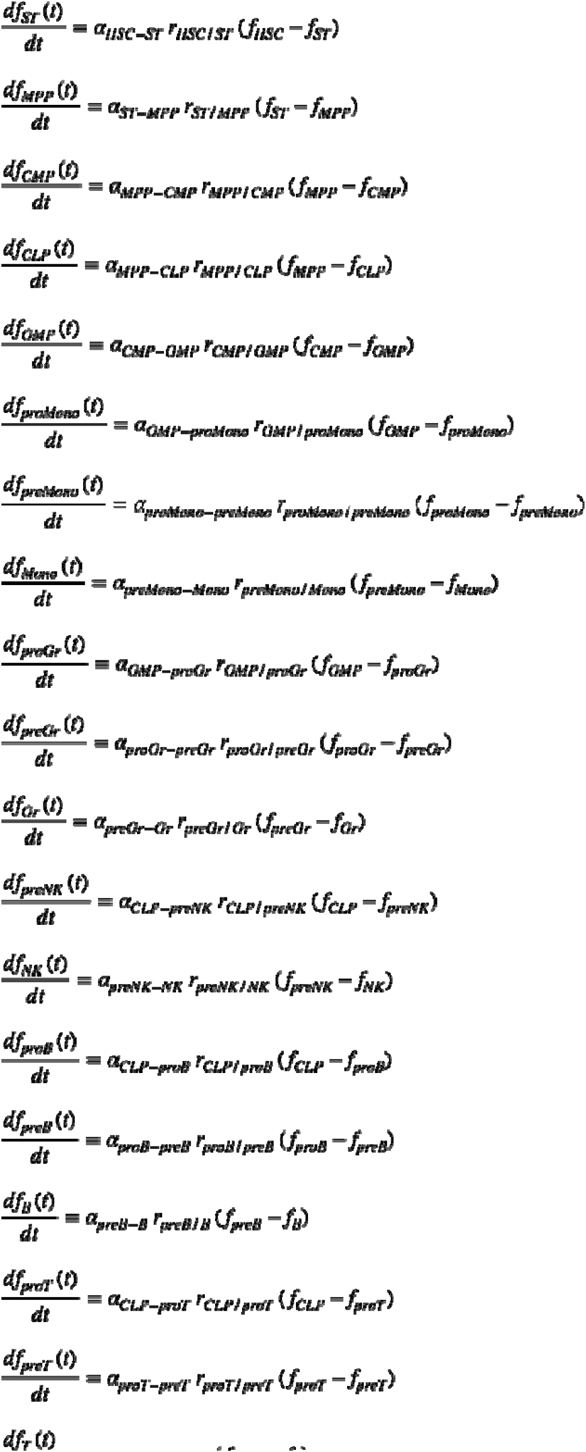

where *f*_*p*_ is the label frequency in population *p*, α_*p1-p2*_ is the differentiation rate of cells from population *p*_*1*_ to population *p*_*2*_ and *r*_*p1/p2*_ is the ratio of population sizes *p*_*1*_ *to p*_*2*_. The ordinary differential equations relate the rate of change of label fraction in a product population to the differentiation rate from its precursor population, as well as the compartment size ratio between the precursor and product populations and can be derived as described by Busch and colleagues^21^. When modeling label dynamics in *Flt3CreERT2tgRosa*^*PolyloxExpress*^ mice, initial label frequencies were estimated together with differentiation rates for stem and progenitor compartments down to CMP and CLP and were set to zero for all downstream compartments. Differentiation rates were inferred by fitting label dynamics in mature blood populations in P6-labeled *Flt3CreERT2tgRosa*^*PolyloxExpress*^ animals and as well as label dynamics in ST and MPP compartments in adult-labeled *Tie2*^*MCM/+*^*Rosa*^*YFP*^ mice^21^. Population size ratios among stem and progenitor populations, as well as GMP, CLP and mature cells were determined based on available data^19,21^ and were assumed to increase linearly through the maturation stages from GMP and CLP towards mature populations. Differentiation rate estimates were computed using Turing.jl, a Bayesian inference library. Maturation times were calculated by summing the reciprocals of the median differentiation rates over all differentiation steps from stem or progenitor compartment to the respective product compartment.

### Stochastic simulations

The following system was simulated stochastically to produce barcode sharing patterns shown in Fig. S3E.

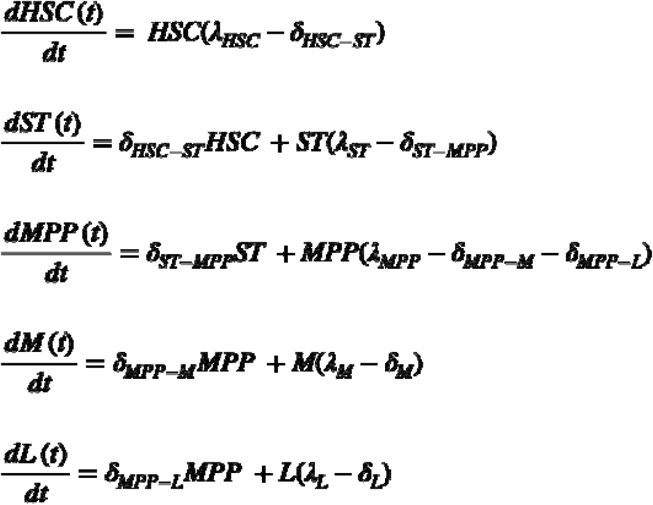

Forward simulation was performed using the Gillespie algorithm and the following rates (given in d^-1^): λ_HSC_=0.01, δ_HSC-ST_=0.01, λ_ST_=0.05, δ_ST-MPP_=0.051, λ_MPP_=0.8, δ_MPP-M_=0.61, δ_MPP-L_=0.41, λ_M_=1.5, δ_M_=1.6, λ_L_=1.0, δ_L_=1.1. Simulations were initialized with a single cell starting in either HSC or MPP compartments, corresponding to HSC or MPP labeling, respectively and run until day 100.

## Data and Code Availability

All data and codes used in this study are available upon request. Requests for resources should be directed to Hans-Reimer Rodewald.

## Acknowledgements

We thank all members of the Rodewald laboratory for ongoing support and discussions. Sven Schaefer for support with mouse husbandry.

## Author contributions

B.C. designed, performed and analyzed all experiments with support from C.H-K., T.N. carried out computational post-processing of the barcoding data and modeling analysis. B.C., T.N. and H.R.R. wrote the manuscript. N.D. performed experiments and reviewed and edited the manuscript. H.R.R. and T.H. supervised the study, reviewed and edited the manuscript. The manuscript was approved by all authors.

## Author information

The authors declare no competing interests.

## Data and material availability

All data, codes, and materials used in this study are available upon request. Requests for resources should be directed to H.R.R.

## Figure Legends

**Fig. S1.**
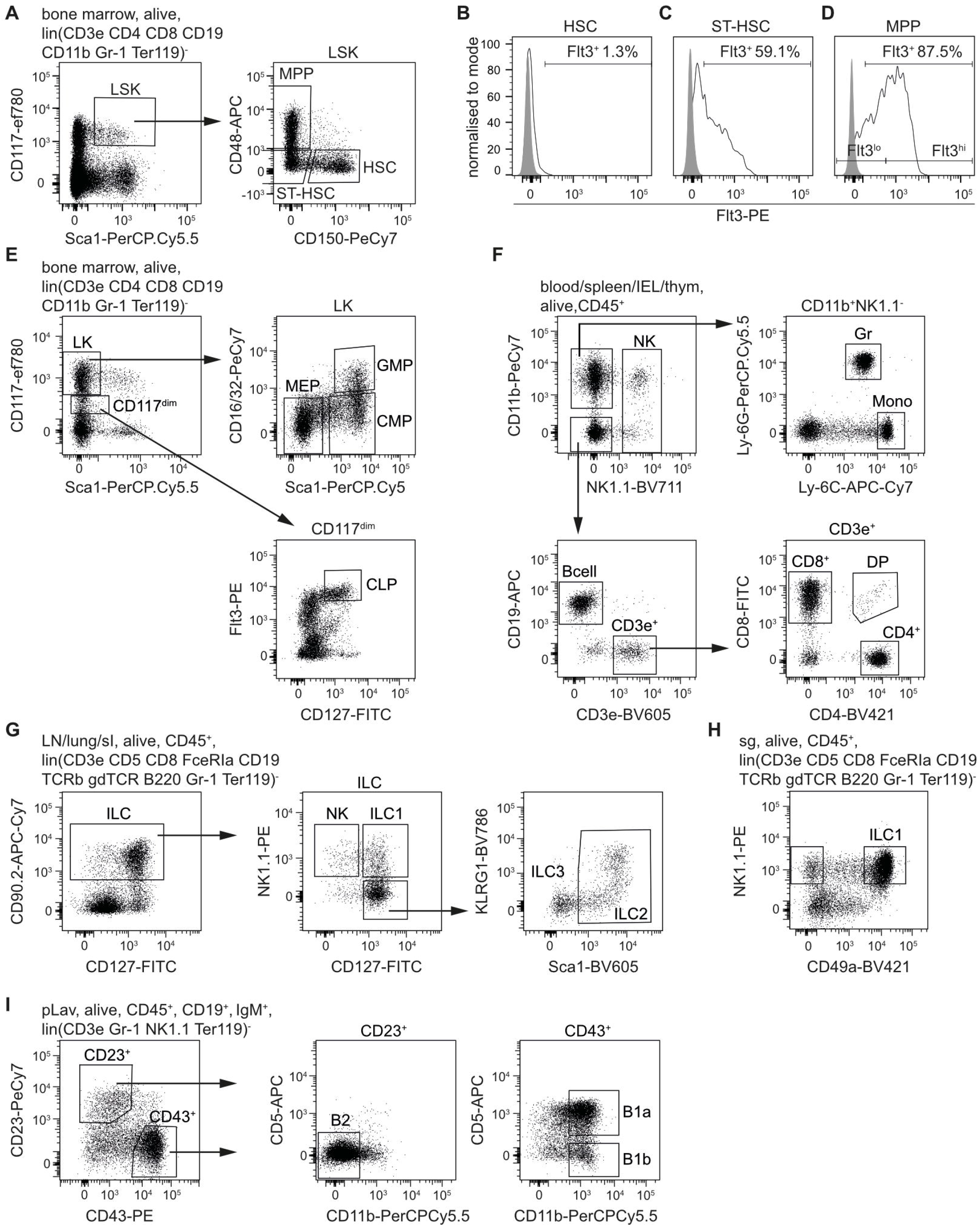
Barcode initiation in early postnatal Flt3^+^ HPC downstream of HSC. A) Gating strategy for early hematopoietic stem and progenitor cells (HSPC) including HSC, ST-HSC and MPP. Flt3 expression in HSC (B), ST-HSC (C) and MPP (D). The “Fluorescence minus one”-control signal is shaded in gray. Flt3^lo^ and Flt3^hi^ gating indicates MPP subpopulations with low and high Flt3 expression, respectively. E) Gating strategy to delineate late HPC including MEP (megakaryocyte-erythrocyte progenitor), CMP (common myeloid progenitor), GMP (granulocyte-monocyte progenitor) and CLP (common lymphoid progenitor) in the bone marrow. F) Gating strategy for the identification of major (circulating) immune cell populations in the blood, spleen, IEL or thymus. G) Gating strategy for the identification of ILC subtypes in lymph nodes, lung or small intestine. H) Gating strategy for the identification ILC1 in the salivary gland (sg). I) Gating strategy for the identification of innate and conventional B cell subtypes from peritoneal lavage cell suspension.

**Fig. S2.**
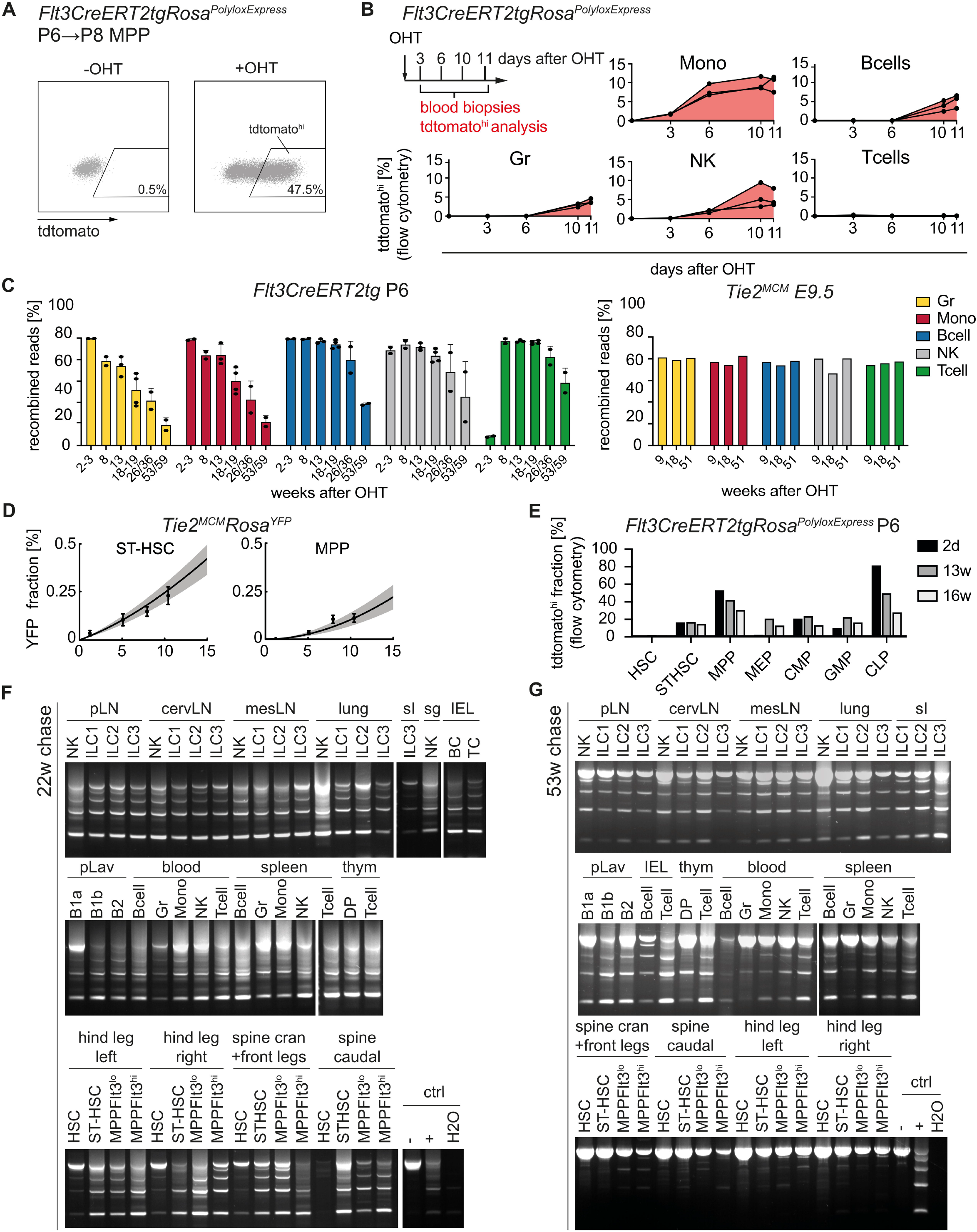
Label propagation dynamics of early postnatal Flt3^+^ HPC. A) Representative flow cytometric analysis of tdtomato^hi^ induction in MPP two days after OHT administration in P6 *Flt3CreERT2tgRosa*^*PolyloxExpress*^ animals (right panel, “+OHT”). Uninduced control mouse (“-OHT”, left panel). B) Short-term labeling kinetics in adult *Flt3CreERT2tgRosa*^*PolyloxExpress*^ OHT-induced animals. The fraction of tdtomato^hi^ cells within major myeloid and lymphoid lineages was assessed within the chasing period of 11 days. Serial bleedings were performed from 3 mice in total. C) Fraction of barcoded circulating cells based on the percentage of reads in non-germline barcode configuration (“recombined reads”) in *Flt3CreERT2tgRosa*^*Polylox*^ animals induced at P6 and analysed over the course of 59 weeks (left) or *Tie2*^*MCM*^*Rosa*^*Polylox*^ analysed over the course of 51 weeks. D) Model fit of label propagation in ST-HSC and MPP in *Tie2*^*MCM*^*Rosa*^*YFP*^ mice^21^. E) Fraction of tdtomato^hi^ cells in HSPC from *Flt3CreERT2tgRosa*^*PolyloxExpress*^ OHT-induced mice that were analysed after 2 days, 13 weeks or 16 weeks (n=1 per end point). F) Amplification of the barcode cassette of enriched cell populations from *Flt3CreERT2tgRosa*^*Polylox*^ animals injected once with OHT at P6 and analysed 22 weeks after induction. pLN, peripheral lymph nodes; cervLN, cercival lymph nodes; mesLN, mesenteric lymph nodes; pLav, peritoneal lavage; sI, small intestine; sg, salivary gland; IEL, intraepithelial cells of the small intestine. G) As in (G) for end point analysis 53 weeks after barcode induction.

**Fig. S3.**
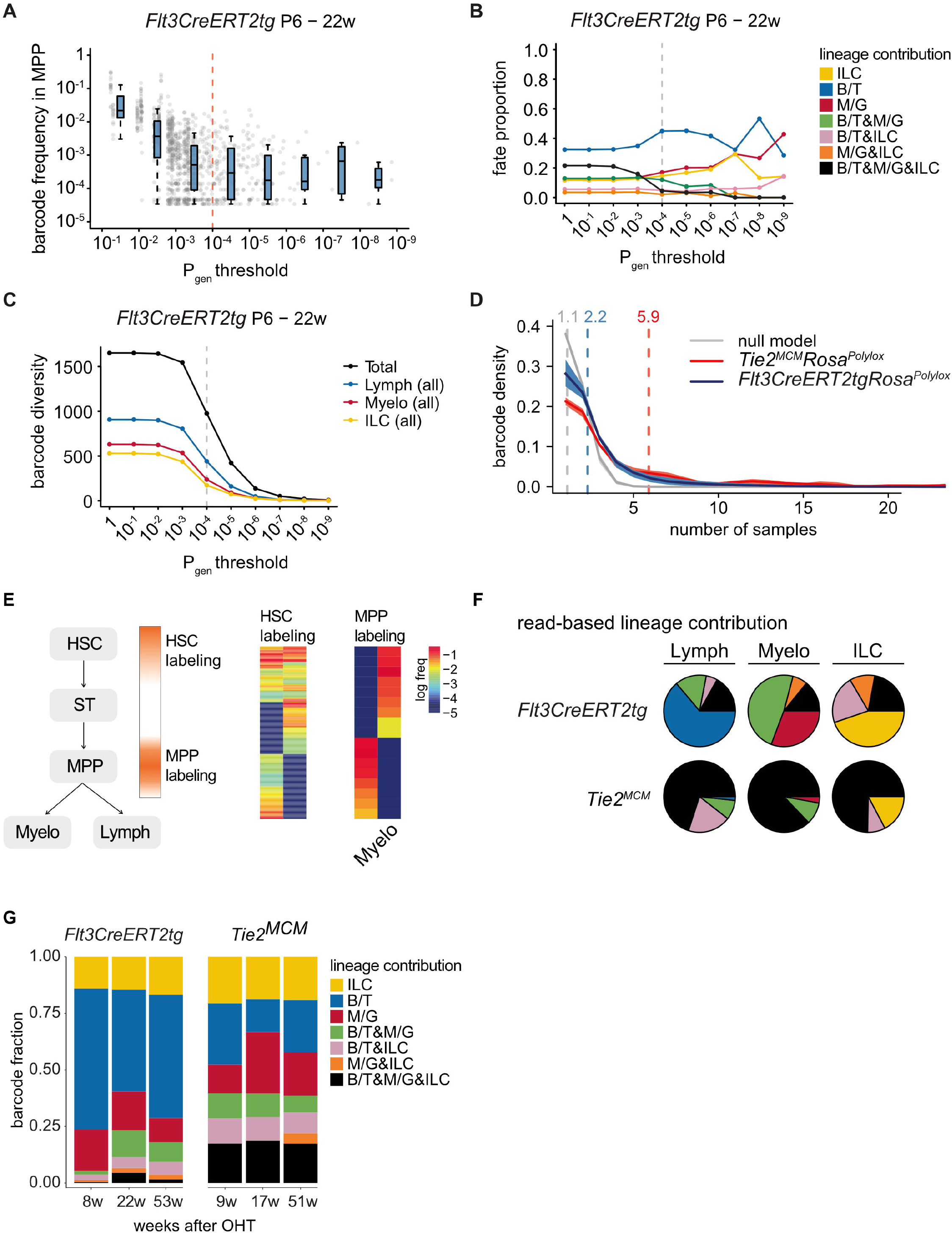
Clonal Flt3^+^ HPC fate combination towards major leukocyte lineages. A) A representative barcode frequency decorrelation analysis on barcodes observed in a P6-induced *Flt3CreERT2tgRosa*^*Polylox*^ mouse. Observed barcodes are binned based on their *P*_gen_ and barcode frequency in MPP samples is shown for each bin. At large *P*_gen_, barcode frequency is correlated with *P*_gen_. However, the correlation is no longer observed for barcodes with *P*_gen_ < 10^-4^. This effect is due to barcode induction of highly probable barcodes in several cells, while barcodes with low generation probability are induced in the smallest possible unit, a single cell. The decorrelation point, *P*_gen_ = 10^-4^, thus marks the threshold for clonal barcodes. B) Proportion of unilineage, bilineage and multilineage fates is depicted for different *P*_gen_ thresholds for a representative *Flt3CreERT2tgRosa*^*Polylox*^ mouse. Dashed vertical line marks the threshold used for classification of clonal barcodes. C) Number of observed barcodes (barcode diversity) in a representative *Flt3CreERT2tgRosa*^*Polylox*^ mouse is depicted for different *P*_gen_ thresholds. Total number of barcodes in the mouse is shown in black and contains barcodes observed in all populations and locations. Number of barcodes in B and T cell populations across all locations are shown in blue (Lymph) and Monocytes and Granulocytes in red (Myelo). Barcodes observed in all ILC populations are shown in yellow. Dashed vertical line marks the threshold used for classification of clonal barcodes. D) Quantification of barcode sharing among samples collected from individual mice. Distributions show the fraction of clonal barcodes (*P*_gen_ < 10E-4) being shared by a certain number of samples from a mouse. At least 50 samples were taken from each mouse and spanned multiple blood populations and organs. Average distributions for *Tie2*^*MCM*^*-* (n=3) and *Flt3CreERT2tg-*labeled (n=7) mice up to 20 samples are shown in red and blue, respectively. The simulated distribution for the null model (grey) was obtained by independently barcoding populations *in silico* using the barcode generation probability model (Pei et. al, 2017) until barcode numbers were matching the experiments. The average number of populations in which a barcode is found is marked with a dashed line and the number on top indicates distribution average. While clonal barcodes are found in several populations on average in *Tie2*^*MCM*^*Rosa*^*Polylox*^ and *Flt3CreERT2tgRosa*^*Polylox*^ mice, they are rarely shared in unrelated populations. E) Effect of barcode induction at different progenitor stages on barcode sharing among myeloid and lymphoid populations. Stochastic simulations were performed starting with a single barcoded HSC or MPP. The cells could proliferate and differentiate into either myeloid or the lymphoid lineage. Heatmaps on the right show the frequency of barcoded progeny in the two lineages for these two labeling scenarios. A significant proportion of single HSC progeny can appear in both lineages (barcode sharing), while barcoded progeny of single MPPs only appear in one lineage (unique barcodes). F) Read-count based quantification of barcode sharing between lineages in *Flt3CreERT2tgRosa*^*Polylox*^ (top) and *Tie2*^*MCM*^*Rosa*^*Polylox*^ (bottom). Related to Fig. 3D. G) Sharing comparison of barcodes initiated in three *Flt3CreERT2tgRosa*^*Polylox*^ animals at P6 (left) and three *Tie2*^*MCM*^*Rosa*^*Polylox*^ animals at E9.5 (right) sacrificed and analyzed at the indicated timepoints. Fraction of barcodes overlapping in different lineages are color-coded.

**Fig. S4.**
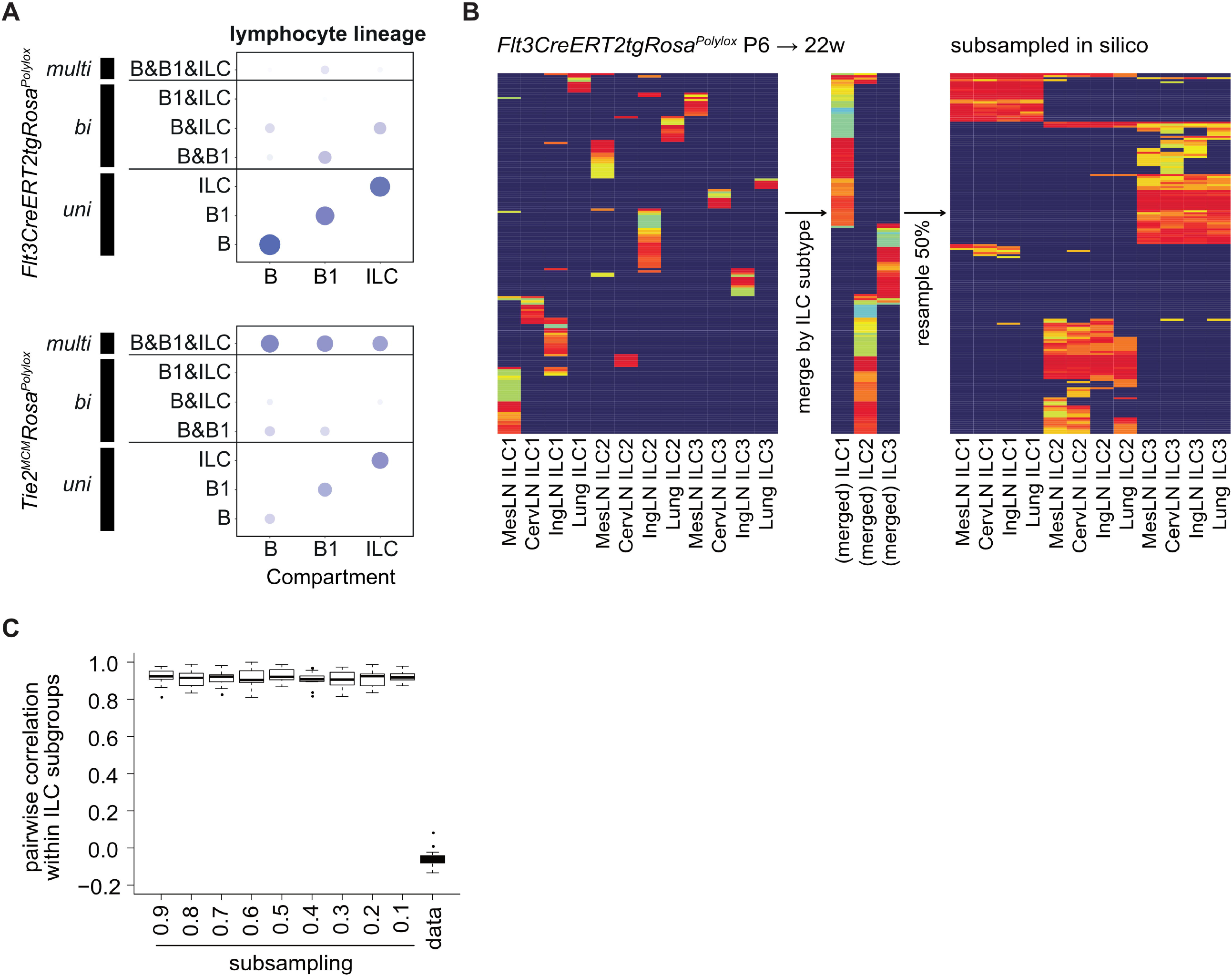
Subset- and tissue-resolved fate specification of P6-labelled Flt3^+^ HPC clones. A) Lymphocyte-centric bubble plots including barcodes detected in conventional B cells, B1 cells and ILC. Co-occurrence of barcodes in additional compartments is indicated by bubble size and shading. Related to Figure 4A. B) Heatmap of ILC1, ILC2 and ILC3 sampled from four different sites at 22 weeks post induction in a P6-induced *Flt3CreERT2tg*^*RosaPolylox*^ mouse (left) and a heatmap for merged ILC1, ILC2 and ILC3 populations from the same mouse, where barcode frequencies from four sites were averaged (middle). *In silico* barcode sampling at a 50% subsampling rate from the merged ILC populations (right). C) Effects of subsampling on the correlation of barcode frequencies among ILC samples (related to Figure 4SB). Progressive subsampling from the merged barcode frequencies was performed from 90% to 10% of experimental read counts for the respective ILC populations. Low correlations in the data could not be achieved by subsampling alone.

## References

1. Orkin, S.H., and Zon, L.I. (2008). Hematopoiesis: an evolving paradigm for stem cell biology. Cell 132, 631–644. 10.1016/j.cell.2008.01.025.

2. Laurenti, E., and Gottgens, B. (2018). From haematopoietic stem cells to complex differentiation landscapes. Nature 553, 418–426. 10.1038/nature25022.

3. Rodriguez-Fraticelli, A.E. (2025). Clonal tracing of blood stem cells across mouse and human lifespans. Blood. 10.1182/blood.2024028195.

4. Hofer, T., Busch, K., Klapproth, K., and Rodewald, H.R. (2016). Fate Mapping and Quantitation of Hematopoiesis In Vivo. Annu Rev Immunol 34, 449–478. 10.1146/annurev-immunol-032414-112019.

5. Christodoulou, C., Spencer, J.A., Yeh, S.A., Turcotte, R., Kokkaliaris, K.D., Panero, R., Ramos, A., Guo, G., Seyedhassantehrani, N., Esipova, T.V., et al. (2020). Live-animal imaging of native haematopoietic stem and progenitor cells. Nature 578, 278–283. 10.1038/s41586-020-1971-z.

6. Christensen, J.L., and Weissman, I.L. (2001). Flk-2 is a marker in hematopoietic stem cell differentiation: a simple method to isolate long-term stem cells. Proc Natl Acad Sci U S A 98, 14541–14546. 10.1073/pnas.261562798.

7. Boyer, S.W., Schroeder, A.V., Smith-Berdan, S., and Forsberg, E.C. (2011). All hematopoietic cells develop from hematopoietic stem cells through Flk2/Flt3-positive progenitor cells. Cell Stem Cell 9, 64–73. 10.1016/j.stem.2011.04.021.

8. Adolfsson, J., Borge, O.J., Bryder, D., Theilgaard-Monch, K., Astrand-Grundstrom, I., Sitnicka, E., Sasaki, Y., and Jacobsen, S.E. (2001). Upregulation of Flt3 expression within the bone marrow Lin(-)Sca1(+)c-kit(+) stem cell compartment is accompanied by loss of self-renewal capacity. Immunity 15, 659–669. 10.1016/s1074-7613(01)00220-5.

9. Fanti, A.K., Busch, K., Greco, A., Wang, X., Cirovic, B., Shang, F., Nizharadze, T., Frank, L., Barile, M., Feyerabend, T.B., et al. (2023). Flt3- and Tie2-Cre tracing identifies regeneration in sepsis from multipotent progenitors but not hematopoietic stem cells. Cell Stem Cell 30, 207–218 e207. 10.1016/j.stem.2022.12.014.

10. Munz, C.M., Dressel, N., Chen, M., Grinenko, T., Roers, A., and Gerbaulet, A. (2023). Regeneration following blood loss and acute inflammation proceeds without contribution of primitive HSCs. Blood. 10.1182/blood.2022018996.

11. Sommerkamp, P., Romero-Mulero, M.C., Narr, A., Ladel, L., Hustin, L.S.P., Schonberger, K., Renders, S., Altamura, S., Zeisberger, P., Jacklein, K., et al. (2021). Mouse multipotent progenitor 5 cells are located at the interphase between hematopoietic stem and progenitor cells. Blood. 10.1182/blood.2020007876.

12. Patel, S.H., Christodoulou, C., Weinreb, C., Yu, Q., da Rocha, E.L., Pepe-Mooney, B.J., Bowling, S., Li, L., Osorio, F.G., Daley, G.Q., and Camargo, F.D. (2022). Lifelong multilineage contribution by embryonic-born blood progenitors. Nature 606, 747–753. 10.1038/s41586-022-04804-z.

13. Pei, W., Feyerabend, T.B., Rossler, J., Wang, X., Postrach, D., Busch, K., Rode, I., Klapproth, K., Dietlein, N., Quedenau, C., et al. (2017). Polylox barcoding reveals haematopoietic stem cell fates realized in vivo. Nature 548, 456–460. 10.1038/nature23653.

14. Pei, W., Wang, X., Rossler, J., Feyerabend, T.B., Hofer, T., and Rodewald, H.R. (2019). Using Cre-recombinase-driven Polylox barcoding for in vivo fate mapping in mice. Nat Protoc. 10.1038/s41596-019-0163-5.

15. Benz, C., Martins, V.C., Radtke, F., and Bleul, C.C. (2008). The stream of precursors that colonizes the thymus proceeds selectively through the early T lineage precursor stage of T cell development. J Exp Med 205, 1187–1199. 10.1084/jem.20072168.

16. Henneke, P., Kierdorf, K., Hall, L.J., Sperandio, M., and Hornef, M. (2021). Perinatal development of innate immune topology. Elife 10. 10.7554/eLife.67793.

17. Torow, N., Hand, T.W., and Hornef, M.W. (2023). Programmed and environmental determinants driving neonatal mucosal immune development. Immunity 56, 485–499. 10.1016/j.immuni.2023.02.013.

18. Pei, W., Shang, F., Wang, X., Fanti, A.K., Greco, A., Busch, K., Klapproth, K., Zhang, Q., Quedenau, C., Sauer, S., et al. (2020). Resolving Fates and Single-Cell Transcriptomes of Hematopoietic Stem Cell Clones by PolyloxExpress Barcoding. Cell Stem Cell 27, 383–395 e388. 10.1016/j.stem.2020.07.018.

19. Cosgrove, J., Hustin, L.S.P., de Boer, R.J., and Perie, L. (2021). Hematopoiesis in numbers. Trends Immunol 42, 1100–1112. 10.1016/j.it.2021.10.006.

20. Hofer, T., and Rodewald, H.R. (2016). Output without input: the lifelong productivity of hematopoietic stem cells. Curr Opin Cell Biol 43, 69–77. 10.1016/j.ceb.2016.08.003.

21. Busch, K., Klapproth, K., Barile, M., Flossdorf, M., Holland-Letz, T., Schlenner, S.M., Reth, M., Hofer, T., and Rodewald, H.R. (2015). Fundamental properties of unperturbed haematopoiesis from stem cells in vivo. Nature 518, 542–546. 10.1038/nature14242.

22. Nizharadze, T., Busch, K., Fanti, A.K., Rodewald, H.R., and Hofer, T. (2023). Differentiation tracing identifies hematopoietic regeneration from multipotent progenitors but not stem cells. Cells Dev 175, 203861. 10.1016/j.cdev.2023.203861.

23. Hofer, T., Barile, M., and Flossdorf, M. (2016). Stem-cell dynamics and lineage topology from in vivo fate mapping in the hematopoietic system. Curr Opin Biotechnol 39, 150–156. 10.1016/j.copbio.2016.04.001.

24. Shang, F., Nizharadze, T., Thiele, R., Cirovic, B., Frank, L., Busch, K., Pei, W., Feyerabend, T.B., Hofer, T., Wang, X., and Rodewald, H.R. (2025). Multipotent progenitors with distinct origins, clonal lineage fates, transcriptomes, and surface markers yield two hematopoietic trees. Proc Natl Acad Sci U S A 122, e2505510122. 10.1073/pnas.2505510122.

25. Sun, J., Ramos, A., Chapman, B., Johnnidis, J.B., Le, L., Ho, Y.J., Klein, A., Hofmann, O., and Camargo, F.D. (2014). Clonal dynamics of native haematopoiesis. Nature 514, 322–327. 10.1038/nature13824.

26. Yokomizo, T., Ideue, T., Morino-Koga, S., Tham, C.Y., Sato, T., Takeda, N., Kubota, Y., Kurokawa, M., Komatsu, N., Ogawa, M., et al. (2022). Independent origins of fetal liver haematopoietic stem and progenitor cells. Nature. 10.1038/s41586-022-05203-0.

27. Xu, W., Cherrier, D.E., Chea, S., Vosshenrich, C., Serafini, N., Petit, M., Liu, P., Golub, R., and Di Santo, J.P. (2019). An Id2(RFP)-Reporter Mouse Redefines Innate Lymphoid Cell Precursor Potentials. Immunity. 10.1016/j.immuni.2019.02.022.

28. Harly, C., Kenney, D., Ren, G., Lai, B., Raabe, T., Yang, Q., Cam, M.C., Xue, H.H., Zhao, K., and Bhandoola, A. (2019). The transcription factor TCF-1 enforces commitment to the innate lymphoid cell lineage. Nat Immunol. 10.1038/s41590-019-0445-7.

29. Kasal, D.N., and Bendelac, A. (2021). Multi-transcription factor reporter mice delineate early precursors to the ILC and LTi lineages. J Exp Med 218. 10.1084/jem.20200487.

30. Walker, J.A., Clark, P.A., Crisp, A., Barlow, J.L., Szeto, A., Ferreira, A.C.F., Rana, B.M.J., Jolin, H.E., Rodriguez-Rodriguez, N., Sivasubramaniam, M., et al. (2019). Polychromic Reporter Mice Reveal Unappreciated Innate Lymphoid Cell Progenitor Heterogeneity and Elusive ILC3 Progenitors in Bone Marrow. Immunity 51, 104–118 e107. 10.1016/j.immuni.2019.05.002.

31. Gasteiger, G., Fan, X., Dikiy, S., Lee, S.Y., and Rudensky, A.Y. (2015). Tissue residency of innate lymphoid cells in lymphoid and nonlymphoid organs. Science 350, 981–985. 10.1126/science.aac9593.

32. Zeis, P., Lian, M., Fan, X., Herman, J.S., Hernandez, D.C., Gentek, R., Elias, S., Symowski, C., Knopper, K., Peltokangas, N., et al. (2020). In Situ Maturation and Tissue Adaptation of Type 2 Innate Lymphoid Cell Progenitors. Immunity 53, 775–792 e779. 10.1016/j.immuni.2020.09.002.

33. Bai, L., Vienne, M., Tang, L., Kerdiles, Y., Etiennot, M., Escaliere, B., Galluso, J., Wei, H., Sun, R., Vivier, E., et al. (2021). Liver type 1 innate lymphoid cells develop locally via an interferon-gamma-dependent loop. Science 371. 10.1126/science.aba4177.

34. Cherrier, D.E., Serafini, N., and Di Santo, J.P. (2018). Innate Lymphoid Cell Development: A T Cell Perspective. Immunity 48, 1091–1103. 10.1016/j.immuni.2018.05.010.

35. Harly, C., Cam, M., Kaye, J., and Bhandoola, A. (2018). Development and differentiation of early innate lymphoid progenitors. J Exp Med 215, 249–262. 10.1084/jem.20170832.

36. Huang, Q., Seillet, C., and Belz, G.T. (2017). Shaping Innate Lymphoid Cell Diversity. Front Immunol 8, 1569. 10.3389/fimmu.2017.01569.

37. Kobayashi, M., Wei, H., Yamanashi, T., Azevedo Portilho, N., Cornelius, S., Valiente, N., Nishida, C., Cheng, H., Latorre, A., Zheng, W.J., et al. (2023). HSC-independent definitive hematopoiesis persists into adult life. Cell Rep 42, 112239. 10.1016/j.celrep.2023.112239.

