## Supplemental Information 1 for "Early postnatal Flt3^+^ hematopoietic progenitors realize fate-restricted and long-lived output *in vivo*"

| Primer ID | Primer Sequence [5'-3'] |
|---|---|
| FW_2426_A | GGTAGTCTCCATCCGACGACACTGCCAAAGATTTC |
| FW_2426_B | GGTAGTCTGGTTGCGACGACACTGCCAAAGATTTC |
| FW_2426_C | GGTAGTCCGACAACGACGACACTGCCAAAGATTTC |
| FW_2426_D | GGTAGTCACGAACCGACGACACTGCCAAAGATTTC |
| FW_2426_E | GGTAGTACGGTACCGACGACACTGCCAAAGATTTC |
| FW_2426_F | GGTAGTAGCCAAGCGACGACACTGCCAAAGATTTC |
| FW_2426_G | GGTAGCTCTTCGACGACGACACTGCCAAAGATTTC |
| FW_2426_H | GGTAGCCTTAGACCGACGACACTGCCAAAGATTTC |
| FW_2426_I | GGTAGAAGGTCCTCGACGACACTGCCAAAGATTTC |
| FW_2426_J | GGTAGTTCGAGGTCGACGACACTGCCAAAGATTTC |
| FW_2426_K | GGTAGTTCACCGACGACGACACTGCCAAAGATTTC |
| FW_2426_L | GGTAGTTGTGAGGCGACGACACTGCCAAAGATTTC |
| FW_2426_M | GGTAGAGGTTGCACGACGACACTGCCAAAGATTTC |
| FW_2426_N | GGTAGTTGCTGTCCGACGACACTGCCAAAGATTTC |
| FW_2426_O | GGTAGCATCATGGCGACGACACTGCCAAAGATTTC |
| FW_2426_P | GGTAGGCATCCTTCGACGACACTGCCAAAGATTTC |
| FW_2426_Q | GGTAGGGTCTTGTCGACGACACTGCCAAAGATTTC |
| FW_2426_R | GGTAGAGCTTCACCGACGACACTGCCAAAGATTTC |
| FW_2426_S | GGTAGAACGCACTCGACGACACTGCCAAAGATTTC |
| FW_2426_T | GGTAGAAGTAGCCCGACGACACTGCCAAAGATTTC |
| FW_2426_U | GGTAGTTGCTCCACGACGACACTGCCAAAGATTTC |
| FW_2426_V | GGTAGGGAAGTGACGACGACACTGCCAAAGATTTC |
| FW_2426_W | GGTAGGACAAGTGCGACGACACTGCCAAAGATTTC |
| FW_2426_X | GGTAGAATAGCGGCGACGACACTGCCAAAGATTTC |
| RV_2427_1 | GGTAGAGACCAAGCATACCTTAGAGAAAGCCTGTCGAG |
| RV_2427_2 | GGTAGGAAGTAGGCATACCTTAGAGAAAGCCTGTCGAG |
| RV_2427_3 | GGTAGATAGACGGCATACCTTAGAGAAAGCCTGTCGAG |
| RV_2427_4 | GGTAGGAACGGAACATACCTTAGAGAAAGCCTGTCGAG |
| RV_2427_5 | GGTAGGACATTGGCATACCTTAGAGAAAGCCTGTCGAG |
| RV_2427_6 | GGTAGTGATCACGCATACCTTAGAGAAAGCCTGTCGAG |
| RV_2427_7 | GGTAGAACATCCGCATACCTTAGAGAAAGCCTGTCGAG |
| RV_2427_8 | GGTAGAATCGTCGCATACCTTAGAGAAAGCCTGTCGAG |
| RV_2427_9 | GGTAGTTAGGCTGCATACCTTAGAGAAAGCCTGTCGAG |
| RV_2427_10 | GGTAGTGGTCTTGCATACCTTAGAGAAAGCCTGTCGAG |
| RV_2427_11 | GGTAGCATTCGACCATACCTTAGAGAAAGCCTGTCGAG |
| RV_2427_12 | GGTAGCACACGAACATACCTTAGAGAAAGCCTGTCGAG |
| RV_2427_13 | GGTAGTGTTGACCCATACCTTAGAGAAAGCCTGTCGAG |
| RV_2427_14 | GGTAGACGAATCCCATACCTTAGAGAAAGCCTGTCGAG |
| RV_2427_15 | GGTAGCTGATTCCCATACCTTAGAGAAAGCCTGTCGAG |
| RV_2427_16 | GGTAGGCAGTATCCATACCTTAGAGAAAGCCTGTCGAG |
| RV_2427_17 | GGTAGCTAACCTCCATACCTTAGAGAAAGCCTGTCGAG |
| RV_2427_18 | GGTAGGTTGGTTCCATACCTTAGAGAAAGCCTGTCGAG |
| RV_2427_19 | GGTAGCGTTGCATCATACCTTAGAGAAAGCCTGTCGAG |
| RV_2427_20 | GGTAGACGGTTGTCATACCTTAGAGAAAGCCTGTCGAG |
| RV_2427_21 | GGTAGACCTTGCTCATACCTTAGAGAAAGCCTGTCGAG |
| RV_2427_22 | GGTAGTTGCCTCTCATACCTTAGAGAAAGCCTGTCGAG |
| RV_2427_23 | GGTAGCCTGTTCTCATACCTTAGAGAAAGCCTGTCGAG |
| RV_2427_24 | GGTAGCGCCAATTCATACCTTAGAGAAAGCCTGTCGAG |
