## Supplemental Information 2 for "Early postnatal Flt3^+^ hematopoietic progenitors realize fate-restricted and long-lived output *in vivo*"

| Antibody | Identifier |
|---|---|
| Anti-Mouse Ly-6C APC-Cy7 | BD Biosciences Cat# 560596, RRID:AB_1727555 |
| Anti-Mouse IgM FITC | Thermo Fisher Scientific Cat# 11-5890-85, RRID:AB_465293 |
| Anti-Mouse CD19 BV605 | BioLegend Cat# 115540, RRID:AB_2563067 |
| Anti-Mouse CD43 APC/Fire | BioLegend Cat# 143216, RRID:AB_2800663 |
| Anti-Mouse CD5 APC | Thermo Fisher Scientific Cat# 17-0051-81, RRID:AB_469330 |
| Anti-Mouse CD11b PerCP-Cy5.5 | Thermo Fisher Scientific Cat# 45-0112-82, RRID:AB_953558 |
| Anti-Mouse CD23 PE-Cy7 | Thermo Fisher Scientific Cat# 25-0232-82, RRID:AB_469604 |
| Anti-Mouse CD45 AF700 | Thermo Fisher Scientific Cat# 56-0451-82, RRID:AB_891454 |
| Anti-Mouse NK1.1 BV421 | BioLegend Cat# 108731, RRID:AB_10895916 |
| Anti-Mouse Ly-6G/Ly-6C (Gr-1) BV421 | BioLegend Cat# 108445, RRID:AB_2562903 |
| Anti-Mouse TER-119 BV421 | BioLegend Cat# 116234, RRID:AB_2562917 |
| Anti-Mouse CD3e BV421 | BioLegend Cat# 100341, RRID:AB_2562556 |
| Anti-Mouse CD43 PE | BioLegend Cat# 143206, RRID:AB_11124719 |
| Anti-Mouse CD19 APC | BD Biosciences Cat# 550992, RRID:AB_398483 |
| Anti-Mouse CD11b PE-Cy7 | Thermo Fisher Scientific Cat# 25-0112-82, RRID:AB_469588 |
| Anti-Mouse CD4 BV421 | BioLegend Cat# 100443, RRID:AB_2562557 |
| Anti-Mouse Ly-6G PerCP-Cy5.5 | BD Biosciences Cat# 560602, RRID:AB_1727563 |
| Anti-Mouse NK1.1 BV711 | BioLegend Cat# 108745, RRID:AB_2563286 |
| Anti-Mouse CD3e BV605 | BioLegend Cat# 100351, RRID:AB_2565842 |
| Anti-Mouse Ly-6G/Ly-6C (Gr-1) FITC | BD Biosciences Cat# 553126, RRID:AB_394642 |
| Anti-Mouse CD8a BV421 | BioLegend Cat# 100753, RRID:AB_2562558 |
| Anti-Mouse CD8a FITC | BD Biosciences Cat# 553030, RRID:AB_394568 |
| Anti-Mouse Ly-6A/E (Sca-1) PerCP-Cy5.5 | Abcam Cat# ab93538, RRID:AB_10674912 |
| Anti-Mouse CD8a PE | BD Biosciences Cat# 553032, RRID:AB_394570 |
| Anti-Mouse CD127 FITC | Thermo Fisher Scientific Cat# 11-1271-82, RRID:AB_465195 |
| Anti-Mouse NK1.1 PE | BD Biosciences Cat# 557391, RRID:AB_396674 |
| Anti-Mouse CD90.2 APC-Cy7 | BioLegend Cat# 105328, RRID:AB_10613293 |
| Anti-Mouse KLRG1 BV785 | BioLegend Cat# 138429, RRID:AB_2629749 |
| Anti-Mouse CD11c BV421 | BioLegend Cat# 101236, RRID:AB_11203704 |
| Anti-Mouse Ly-6A/E (Sca-1) BV605 | BioLegend Cat# 108133, RRID:AB_2562275 |
| Anti-Mouse CD8a APC | Thermo Fisher Scientific Cat# 17-0081-83, RRID:AB_469336 |
| Anti-Mouse TCR β chain APC | BioLegend Cat# 109212, RRID:AB_313435 |
| Anti-Mouse TCR gamma/delta APC | Thermo Fisher Scientific Cat# 17-5711-82, RRID:AB_842756 |
| Anti-Mouse FcεRIα APC | BioLegend Cat# 134316, RRID:AB_10640121 |
| Anti-Mouse TER-119 APC | BD Biosciences Cat# 557909, RRID:AB_398635 |
| Anti-Mouse CD3 APC | Thermo Fisher Scientific Cat# 17-0032-82, RRID:AB_10597589 |
| Anti-Mouse CD45R/B220 APC | BD Biosciences Cat# 553092, RRID:AB_398531 |
| Anti-Mouse Ly-6G/Ly-6C (Gr-1) APC | BD Biosciences Cat# 553129, RRID:AB_398532 |
| Anti-Mouse CD49a BV421 | BD Biosciences Cat# 740046, RRID:AB_2739815 |
| Anti-Mouse CD117 eF780 | Thermo Fisher Scientific Cat# 47-1171-82, RRID:AB_1272177 |
| Anti-Mouse CD48 AF700 | BioLegend Cat# 103426, RRID:AB_10612755 |
| Anti-Mouse CD150 PE-Cy7 | BioLegend Cat# 115914, RRID:AB_439797 |
| Anti-Mouse CD135 APC | BioLegend Cat# 135310, RRID:AB_2107050 |
| Anti-Mouse CD34 BV421 | BioLegend Cat# 152208, RRID:AB_2650766 |
| Anti-Mouse CD16/32 PE-Cy7 | Thermo Fisher Scientific Cat# 25-0161-81, RRID:AB_469597 |
| Anti-Mouse CD3e Biotin | Thermo Fisher Scientific Cat# 13-0031-85, RRID:AB_466320 |
| Anti-Mouse TER-119 Biotin | Thermo Fisher Scientific Cat# 13-5921-85, RRID:AB_466798 |
| Anti-Mouse CD11b Biotin | Innovative Research Cat# RM2815, RRID:AB_1464417 |
| Anti-Mouse CD19 Biotin | BD Biosciences Cat# 553784, RRID:AB_395048 |
| Anti-Mouse Ly-6G (Gr-1) Biotin | Thermo Fisher Scientific Cat# 13-5931-85, RRID:AB_466801 |
| Anti-Mouse CD8a Biotin | Thermo Fisher Scientific Cat# 13-0081-85, RRID:AB_466347 |
| Anti-Mouse CD4 Biotin | BD Biosciences Cat# 553728, RRID:AB_395012 |
| Anti-Mouse CD19 PerCP-Cy5.5 | Thermo Fisher Scientific Cat# 45-0193-80, RRID:AB_906215 |
| Anti-Mouse CD11c APC | BioLegend Cat# 117310, RRID:AB_313779 |
